# Integrating Nanopore MinION Sequencing into National Animal Health AMR Surveillance Programs: An Indonesian Pilot Study of Chicken Slaughterhouse Effluent and Rivers

**DOI:** 10.1101/2024.11.14.623636

**Authors:** Rallya Telussa, Puji Rahayu, Thufeil Yunindika, Curtis Kapsak, Kanti Puji Rahayu, Oli Susanti, Imron Suandy, Nuraini Triwijayanti, Aji B. Niasono, Syamsul Ma’arif, Hendra Wibawa, Lestari, Gunawan B. Utomo, Farida C. Zenal, Luuk Schoonman, Lee E. Voth-Gaeddert

## Abstract

Antimicrobial resistance (AMR) poses significant risks to human and animal health while the environment can contribute to its spread. National AMR surveillance programs are pivotal for assessing AMR prevalence, trends, and intervention outcomes, however, integrating advanced surveillance tools can be difficult. This pilot study, conducted by FAO ECTAD Indonesia and DGLAHS, Indonesian Ministry of Agriculture, evaluated the costs and benefits of integrating the Nanopore MinION, Illumina MiSeq, and Sensititre system into a culture-based slaughterhouse-river surveillance system. Water samples were collected from six chicken slaughterhouses and adjacent rivers (pre- and post-treatment effluent, upstream, downstream). Culture-based ESBL and general *E. coli* concentrations were estimated via the WHO Tricycle Protocol, while isolates (n=42) were sequenced (MinION, MiSeq) and antimicrobial susceptibility testing conducted (Sensititre). The Tricycle Protocol results provided estimates of effluent and river concentrations of ESBL and general *E. coli* identifying ESBL-to-general *E. coli* ratios of 13.8% and 6.2%, respectively. Compared to hybrid sequencing assemblies, MinION had a higher concordance than MiSeq for ARG identification (98%), virulence genes (96%), and locations for both (predominately plasmids). Furthermore, MinION concordance with Sensititre AST was 91%. Cost-benefit comparisons suggest sequencing can complement culture-based methods but is dependent on the value placed on the additional information gained.

**Highlights:**

- This study demonstrates the integration of Nanopore MinION sequencing into a national AMR surveillance program, highlighting its potential for monitoring antibiotic-resistant *E. coli* in slaughterhouse effluent and rivers.
- Our findings show that Nanopore MinION sequencing exhibits high concordance with hybrid sequencing assemblies, validating its effectiveness for genomic surveillance in decentralized settings.
- The study evaluates the cost-benefit of using Nanopore MinION, Illumina MiSeq, and Sensititre AST, comparing information gained with cost.

**Graphical Abstract:** 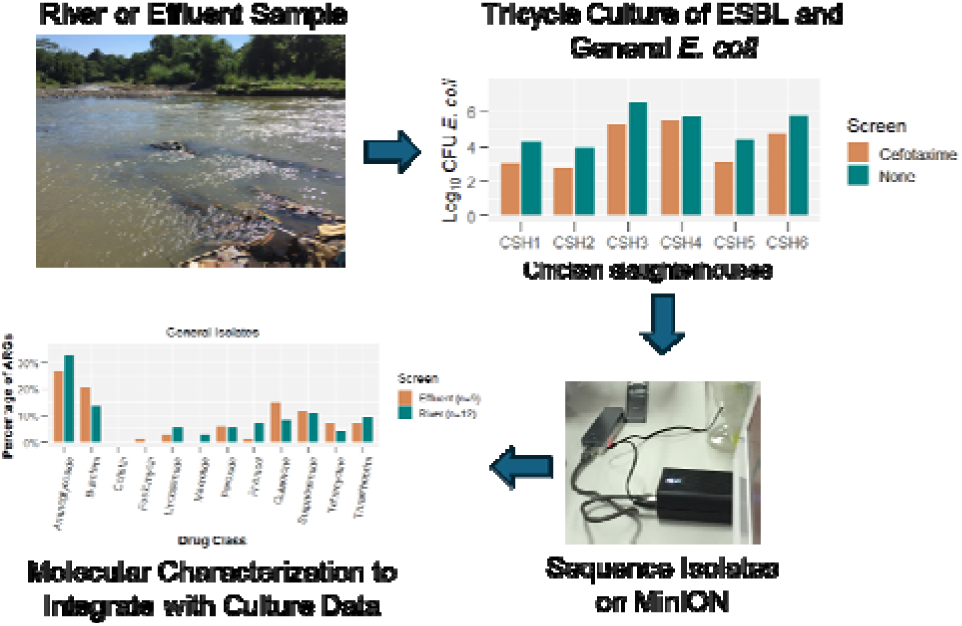

## Introduction

Antimicrobial resistance (AMR) is a critical global health issue. The emergence and spread of AMR is complex and occurs through people, animals, and the environment (Berendonk et al., 2015). To fully understand, monitor, and control the transmission of AMR, a one health approach – people, animals, and the environment – must be leveraged (McEwen and Collignon, 2017). Within this complex one health system (or systems) are critical control points where control strategies can be most effective at curbing AMR transmission (Goulas et al., 2020). Standardized AMR surveillance systems are key for 1) identifying where control strategies would be most effective and 2) measuring the effectiveness of the control strategy. Standardization includes the sampling and laboratory protocols, the types of AMR indicator bacteria evaluated, and the types of environments (and their critical control points) to monitor (Keenum et al., 2022; Liguori et al., 2022; Sano et al., 2020).

Two global AMR surveillance system approaches include 1) the ESBL *Escherichia coli* (*E. coli*) Tricycle Protocol (World Health Organization, 2021, 2016) and 2) PulseNet International (Nadon et al., 2017). The Tricycle Protocol is a proposed, low-cost method that uses a culture-based procedure to estimate the abundance of total *E. coli* and Extended Spectrum Beta Lactamase (ESBL) producing *E. coli* across three sectors (human health, food chain, and the environment). The PulseNet International protocols use a whole genome sequencing (WGS)-based method to evaluate genetic characteristics of individual isolates (clinical and non-clinical). Therefore, understanding the costs and benefits of integrating the two approaches within the context of institutional program operations is important. This can be done by 1) sequencing the isolates from the Tricycle Protocol and 2) validating the use of WGS data for the detection of antimicrobial resistance determinants by comparison to phenotypic Antimicrobial Susceptibility Testing (AST). In addition, understanding differences in results and costs for different sequencing technologies and approaches is important. Comparing outputs from Illumina MiSeq sequencing technology, the Oxford Nanopore MinION sequencing technology, and ‘gold standard’ hybrid assemblies can aid in selecting the optimal technologies and approaches. Finally, bioinformatic tools that do not require command line expertise can allow for further decentralized adoption of these advanced sequencing technologies but require simplified processing and analysis tools.

To evaluate these methods, this pilot study was designed by FAO and DGLAHS with the aim of assessing the concentration and genetic characteristics of ESBL *E. coli* in the effluent and receiving river bodies of chicken slaughterhouses in the Greater Jakarta area (Jakarta, Bogor, Tangerang), Indonesia. Slaughterhouse effluent and rivers are potential locations for monitoring critical control points related to the animal-environment nexus of the one health system for AMR transmission. ESBL-producing *Enterobacteriaceae* is listed as a critical threat to human health by the CDC AR threats report (Centers for Disease Control and Prevention, 2019) supporting the use of ESBL *E. coli* as an indicator bacteria to monitor. Furthermore, recent data suggests chickens and associated slaughterhouses can contribute to the transmission of AMR. Day et al. 2019 suggests ESBL *E. coli* infections among humans are increasing and chickens may contribute to the transmission of pathogenic ESBL *E. coli* between chickens and humans (Day et al., 2019). Further data suggests water (effluent and rivers) can be a primary mode of transmission and spread of ESBL *E. coli* (Blaak et al., 2015; Fuhrmeister et al., 2021; Jorgensen et al., 2017). In Indonesia, the majority of chicken slaughterhouses are located near rivers as there is often a significant level of effluent created with the slaughtering process (personal communication). Furthermore, slaughterhouses are required by law to treat their effluent before discharging into the environment (Directorate of Veterinary Public Health, Ministry of Agriculture, 2021).

In this pilot study, we evaluated the set of methods by sampling and testing slaughterhouse effluent and upstream and downstream river water for six chicken slaughterhouses adjacent to rivers. Questions evaluated in this pilot study included:

1. *what types of E. coli, concentrations, and associated resistance genes are present in the effluent of chicken slaughterhouses and the receiving rivers?*
2. *can the Oxford Nanopore MinION sequencing technology provide valid and valuable data considering costs within a regional Indonesian AMR monitoring systems currently deploying the Tricycle Protocol?*

## Materials & Methods

### Study Site and Sample Collection

The pilot study was conducted in the Greater Jakarta area in Indonesia where a total of six chicken slaughterhouses were sampled. The number of chickens slaughtered per day ranged from 700 to 30,000 per day, with four locations below 4,000 and the remaining two locations above 24,000. Three slaughterhouses operated during the day and three during the night, while three of the six utilized some type of effluent treatment. All slaughterhouse effluent was discharged into adjacent rivers. Samples were collected during operational hours and more than 12 hours after any significant rain event (>2mm over five hours).

Figure 1 depicts the sampling locations at each chicken slaughterhouse (n=6) for the pilot study. At each slaughterhouse, three or four sampling locations were identified. If the slaughterhouse had any type of effluent treatment (n=3), samples were collected before and after treatment (but before entering the river). At all slaughterhouses, river samples were collected both 10 meters upstream and downstream from the discharge point.

**Figure 1.**
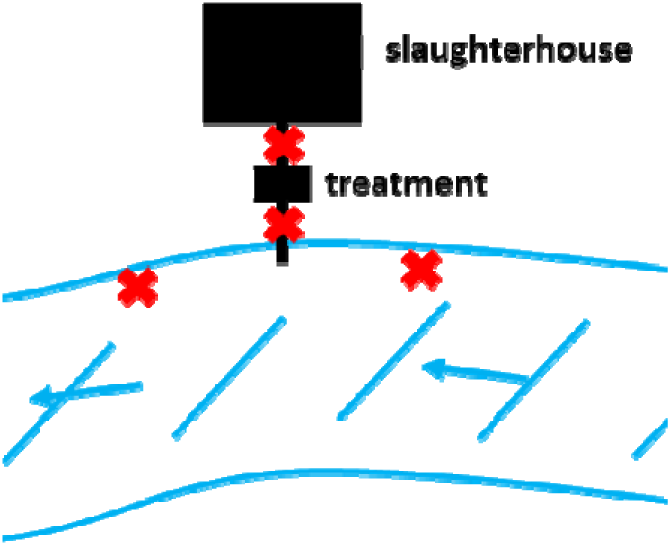
Diagram of sampling points.

For each sampling point, 1000 mL of sample (river water or effluent) was collected utilizing a 300 mL collection beaker and acquiring four samples spaced 1-2 minutes apart and homogenizing the sample via light physical shaking for 30 seconds. Next, the sample was immediately allocated into two sterile glass bottles, a 600 mL bottle and a 250 mL bottle, for direct DNA extraction for metagenomic sequencing and the Tricycle Protocol, respectively. The samples were placed in a cooler and transported back to the Quality Control and Animal Product Certification Laboratory (Balai Pengujian Mutu dan Sertifikasi Produk Hewan; BPMSPH) in Bogor, West Java. In addition, a short survey was used to collect information about the chicken slaughterhouse, the local environment, and the sampling conditions. Additional details of the sampling approach are presented in the supplementary material.

### Tricycle Protocol and Antimicrobial Susceptibility Testing (AST)

On arrival at the laboratory, the 250 mL sample was once again homogenized via light physical shaking for 30 seconds. Enumeration, isolation, and identification of total and ESBL *E. coli* from effluent and river water samples (including ESBL *E. coli* confirmatory tests) were conducted following the membrane filtration version of the Tricycle Protocol (World Health Organization, 2016) (see full workflow in Figure 2 and a detailed protocol in the supplementary material). Briefly, membrane filtration was conducted on a diluted sample (10-fold using PBS) and transferred to a plate with TBX medium for general *E. coli* enumeration (TBX-cefotaxime plate was used for enumeration of presumptive ESBL *E. coli*). Plates were incubated at 37° C for 24 hours. After enumeration of general *E. coli* and presumptive ESBL *E. coli*, 10 general *E. coli* colonies and 10 ESBL *E. coli* colonies were picked from each plate and streaked onto either MacConkey agar plate (general *E. coli*) or MacConkey-cefotaxime plate. Plates were incubated at 37°C for 24 hours. Finally, colonies were picked and transferred to a nutrient agar plate and incubated at 37°C for 12 hours.

**Figure 2.**
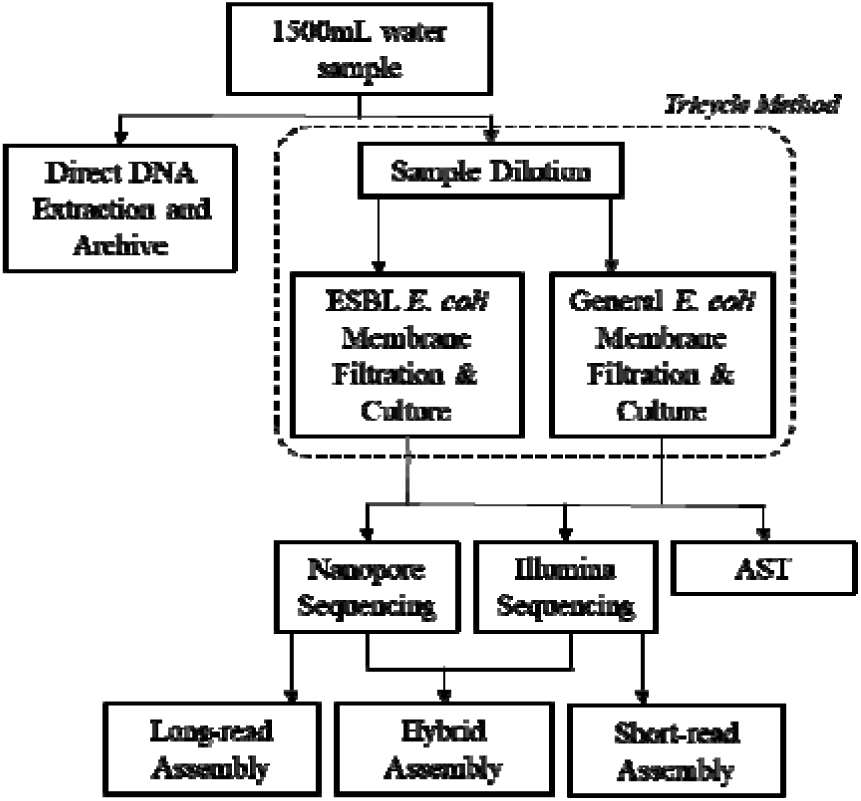
Depiction of the study workflow including Tricycle Protocol for ESBL and general E. coli enumeration, sequencing, and antimicrobial susceptibility testing (AST).

To facilitate identification of *E. coli*, the Sulfide, Indole, Motility (SIM) test, Methyl red and Voges-Proskauer (MRVP) tests, and Citrate tests were conducted (see supplementary material for details). Finally, the double disc method (DDT) method was used to confirm ESBL *E. coli* colonies using cefotaxime, ceftazidime, combination disk of cefotaxime and clavulanic acid, and a combination disk of ceftazidime and clavulanic acid. Bacterial concentrations were then calculated based on the initial enumeration, total volume plated, number of picked colonies, and number of confirmed colonies (we use the term ‘phenotypically confirmed ESBL’ from here on to refer to the presumptive ESBL *E. coli*). Finally, isolates were preserved in glycerol mixed with culture media and stored at −80° C if further testing was not conducted immediately.

Antimicrobial susceptibility testing (AST) for each isolate was conducted using the EU Surveillance Salmonella/*E. coli* (EUVSEC) plate. The Sensititre plate contained 14 different antibiotics (see supplementary material for further details). Following the manufacturer’s protocol, the minimum inhibitory concentrations (MICs) for each antibiotic were determined for all isolates. In this study we define multi-drug resistance (MDR) for phenotypic testing as any isolate that demonstrates an MIC classified as resistant by the Clinical and Laboratory Standards Institute (CLSI) and European Committee on Antimicrobial Susceptibility Testing (EUCAST) breakpoints for at least two antimicrobial agents while for genotypic sequencing any isolate that carries two or more ARGs from different antimicrobial resistance drug classes.

### Whole Genome Sequencing: Oxford Nanopore and Illumina

Whole genome sequencing on the Oxford Nanopore MinION Mk1B (Oxford Nanopore Technologies, Oxford, UK) was conducted at the BPMSPH laboratory. Briefly, DNA extraction of the *E. coli* isolates was conducted using the Qiagen DNeasy Blood and Tissue extraction kit (Hilden, Germany) following the manufacturer’s protocol. DNA quality checks were conducted using the NanoDrop 2000 (ThermoFisher, Waltham, MA) (no Qubit was available). Next, library preparation was conducted using the Rapid Barcoding Sequencing (SQK-RBK004) kit following the manufacturer’s protocol. Initial sequencing runs included only two barcoded isolates per run but increased to four by the last three slaughterhouses (see supplementary material for more information). MinION R9.4 flowcells were used for sequencing.

Whole genome sequencing on the Illumina MiSeq (Illumina, Inc., San Diego, CA, USA) was conducted on a subset of DNA extracts from the *E. coli* isolates at the Disease Investigation Center (DIC) Wates in Yogyakarta, Indonesia. Pure DNA of *E. coli* isolates were transferred on ice to the lab. The PulseNet International protocol for library preparation and sequencing for the Illumina MiSeq was used (Centers for Disease Control and Prevention, 2019b). The Nextera DNA XT kit (Illumina, Inc.) was used for library preparation and sequencing was conducted using the v2 kit with 500 cycles for a read length of 2 x 250 bp.

### Quality Control

Several methods were used to provide quality control and contamination checks throughout the pilot study based on available resources. These included field blanks, deionized water transferred from a storage container to a collection bottle in the field and included in the direct DNA extractions. In addition, deionized water extraction blanks were used for the isolate DNA extractions (post-culture). Quality control strains for the culture-based isolation included *E. coli* 10455 NCSU and Klebsiella pneumoniae ATCC 70 for isolation, identification, and confirmatory tests of ESBL *E. coli*. *E. coli* ATCC 25922 was used as the control strain for isolation and identification of general *E. coli*. Finally, during the initial DNA sequencing of barcoded isolates the lambda control provided in the Nanopore Rapid Barcoding Ligation Kit was included as one of the barcoded samples. Results were validated following the manufacturer’s guidelines.

### Bioinformatic Analyses

For the Nanopore sequencing data, the MinIT device was used to conduct basecalling (using default settings of Guppy v2.2.3) and fast5 and fastq data was transferred to cloud storage and duplicated on an external hard drive. All fastq files were automatically uploaded to Epi2Me (https://epi2me.nanoporetech.com/) for initial contamination and quality control checks. Centrifuge and the NCBI reference database was used to classify reads (Kim et al., 2016). Any sequenced isolate with >5% reads classified as something other than *Escherichia* at the genus level was deemed contaminated.

The fastq files were uploaded to Galaxy Europe (Afgan et al., 2018) for assembly and evaluation. The bioinformatic protocol nanopore Workflow v0.4.4 developed by Katz and Kapsak 2020 (Kapsak and Katz, 2021) was adapted for use on Galaxy Europe (see Figure S1 for pipeline depiction). Briefly, reads were filtered using filtlong v0.2.0 (Wick, 2018) with default parameters and the highest quality reads totaling 600 Mbases were selected for downstream analysis. Quality check (QC) reports were generated with NanoPlot v1.28.2 (Coster et al., 2018). Initial assembly was conducted by Flye v2.3.7 (default settings, estimated genome size 5m) (Kolmogorov et al., 2019) while the filtered fastq files were converted to fasta files (FASTQ to FASTA converter v1.1.5). Next, minimap2 v2.17 (Li, 2018) was used to align reads to the draft assembly from Flye and Racon v1.3.1.1 (Vaser et al., 2017) was used for contig consensus correction. This correction process was repeated four times as recommended by the medaka documentation (Oxford Nanopore Technologies LLC, 2021). Finally, medaka v1.0.1 (Oxford Nanopore Technologies Ltd.) was used as a final consensus correction step before evaluation. PlasFlow v1.0 was used to differentiate between chromosome and plasmid contigs (Krawczyk et al., 2018). Finally, the adapted long-read isolate pipeline on Galaxy Europe was compared with the originally developed long-read isolate pipeline available on Github (https://github.com/kapsakcj/nanoporeWorkflow) by running the same test isolate fastq file on both pipelines. This was also conducted for the adapted short-read pipeline available on Github (https://github.com/lskatz/SneakerNet) using a test short-read fastq file.

For Illumina, the protocol developed by Katz 2020 (Griswold et al., 2021; Katz, 2020) was adapted for use on Galaxy Europe. Briefly, Trimmomatic v0.38 was used to filter reads below Q20 (Bolger et al., 2014). QC reports were generated by FastQC v0.72 and MultiQC v1.9 visualization (Ewels et al., 2016) from raw paired-end reads (independent of R1/R2). Next, Kraken v2.0 (Wood and Salzberg, 2014) was run on the merged paired-end reads (default parameters and standard database). Any sequenced isolate with >5% reads classified as something other than *Escherichia* at the genus level was deemed contaminated. Finally, Shovill v1.1.0 (Seemann, 2020) with the skesa assembler (Souvorov et al., 2018) was used for filtering, assembly, and polishing followed by Prokka v1.14.6 (using Prodigal for gene prediction (Hyatt et al., 2010)) for gene prediction and annotation (Seemann, 2014).

For hybrid assemblies, paired-end reads from Illumina sequencing were filtered via Trimmomatic v0.38 while single long-reads from Nanopore sequencing were filtered via filtlong v0.2.0. SPAdes via Unicycler v0.4.8 (Wick et al., 2017) was used with default parameters to generate hybrid assemblies and PlasFlow v1.0 was used to differentiate contigs associated with the chromosome and potential plasmids.

Final assemblies for all isolates (Nanopore, Illumina, and hybrid assembles) were then uploaded to the following tools on the Center for Genomic Epidemiology website (http://www.genomicepidemiology.org/services/), KmerFinder v3.2 (Bacteria organism) (Larsen et al., 2014), ResFinder v4.1 (Bortolaia et al., 2020) (Escherichia coli*, Assembled Genome/Contigs, 98% identification threshold, 98% minimum length), VirulenceFinder 2.0 (Joensen et al., 2014) (Escherichia coli, 98% identification threshold, 98% minimum length, Assembled or Draft Genome/Contigs), and PlasmidFinder v2.1 (Carattoli et al., 2014) (Enterobacteriaceae, 98% identify threshold, 98% minimum length, Assembled or Draft Genome/Contigs). Isolates were classified by pathotype using the virulence genes present, following the protocols from Franz et al. 2015 (Franz et al., 2015) and Sarowska et al. 2019 (Sarowska et al., 2019). Finally, to generate phylogenetic trees, Roary v3.13.0 (Page et al., 2015) (95% blastp identity, 95% of gene for core classification), RAxML v8.2.4 (Stamatakis, 2014) (default), and iTOL v4 (Letunic and Bork, 2019) were used. Phylogenetic trees were used to visualize isolate clusters and evaluate similarities between sequencing platforms and assembly approaches (Nanopore-only, Illumina-only, and hybrid).

### Analyses

For the Tricycle Protocol data, concentrations and proportions of *E. coli* were estimated. Next, to compare AST-identified phenotypic properties with the associated genes, concordances were calculated for a subset of the isolates (n=21) for the Nanopore MinION. To compare genotypic validity of the Nanopore MinION, results from the identification of ARGs, virulence factors, serotypes, and cgMLST were compared to the ‘gold standard’ hybrid assemblies. In addition, to further evaluate gene annotation and cluster capabilities (e.g., for outbreak investigations), integrated phylogenetic trees were estimated using the method described above for all matched isolates (n=21; Nanopore-only, Illumina-only, and hybrid). Trees were then compared for similar structure and clusters. Finally, costs for the pilot study were used to evaluate a per sample cost, both including and excluding capital costs (i.e. laptop, MinIT device, external hard drive, etc.; see supplementary material for details).

## Results

### E. coli concentrations

The Tricycle Protocol generated results on ESBL and general *E. coli* concentrations for six chicken slaughterhouses (effluent, upstream and downstream). Figure 3 displays the log10 transformed *E. coli* concentrations for 1) ESBL *E. coli*, 2) general *E. coli*, and 3) the ratio between ESBL and general *E. coli*.

**Figure 3.**
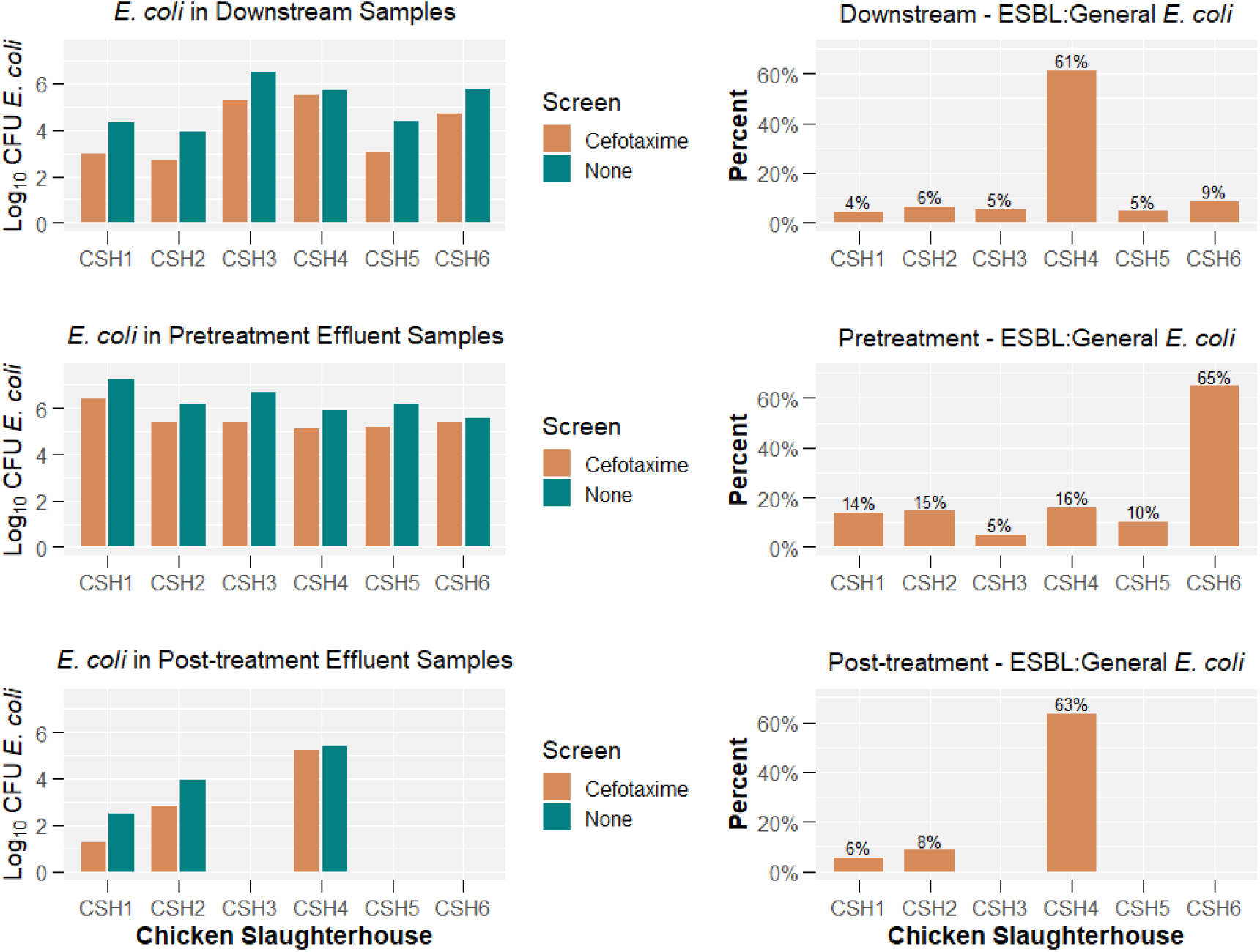
Concentration of ESBL *E. coli* and general *E. coli* across different slaughterhouses and sampling locations.

The median concentrations of ESBL *E. coli* from upstream river samples was 21,244.85 (log10 4.33) CFU/100 mL and from downstream river samples was 24,648.60 (log10 4.39) CFU/100 mL. The median concentrations of ESBL *E. coli* from pre-treatment and post-treatment effluent samples was 229,545.50 (log10 5.36) CFU/100 mL and 709.10 (log10 2.85) CFU/100 mL, respectively. The proportion of ESBL *E. coli* to general *E. coli* in river water samples ranged from 3.5 to 61.3% (mean = 12.2%, median = 6.2%) while effluent samples ranged from 5.0 to 64.7% (mean = 22.4%, median = 13.8%). The proportions varied widely (9 of 12 were between 3.5% −8.6%), however, in effluent direct from the slaughterhouse, the ratio appears to increase (5 of 6 were >10.1%).

Comparing the upstream-downstream change in log10 *E. coli* concentration for both ESBL and general *E. coli* (see Figure 4), seven of 12 measurements suggested the log10 *E. coli* concentration (either general or ESBL) increased downstream as compared to upstream (three decreased and two remained the same). Finally, the data suggested that five of the six measurements in log10 *E. coli* concentration between pre-treatment and post-treatment had pre-treatment concentrations higher than post-treatment (one remained the same).

**Figure 4.**
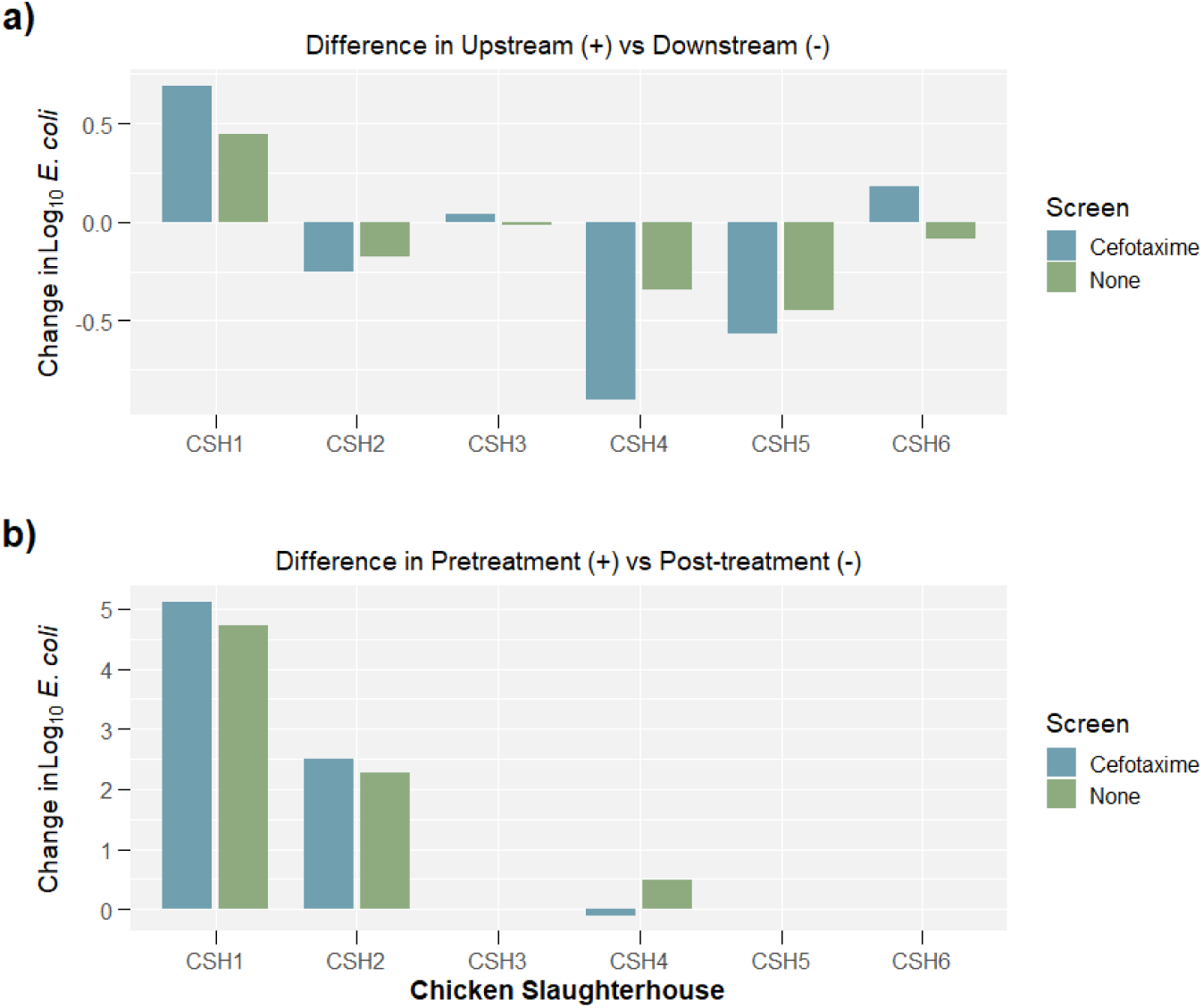
a) Differences in *E. coli* concentrations between upstream and downstream samples and b) pre-treatment and post-treatment samples. *‘+’ denotes that if the concentration was a positive value, there was a higher concentration in ‘Upstream’ or ‘Pretreatment’ samples*.

### Whole Genome Sequencing: Oxford Nanopore vs Illumina

The suggested threshold for the minimum total Mbases per sample (200 Mbases set by PulseNet International) was obtained for all but two samples sequenced on the Oxford Nanopore MinION and all samples sequenced on the Illumina MiSeq. For the Nanopore MinION data, mean read quality post filtering was Q9.9 and per run contamination levels were below the 5% threshold for all isolates. Finally, mean assembly coverage was 96x and the median number of contigs classified as chromosomal was one. For the Illumina MiSeq data, mean read quality post filtering was Q36, per isolate contamination levels were below 5% for all but one isolate (removed for subsequent analyses), mean assembly coverage was 109x and the median number of contigs classified as chromosomal was 52. Table 1 and Table S1 provide sequencing and assembly statistics.

**Table 1.** Sequencing and assembly statistics for Oxford Nanopore (n=42) and Illumina (n=19)

| Results per run | Oxford Nanopore MinION Mean (range) | Illumina MiSeq Mean (range) |
| --- | --- | --- |
| **QC** |  |  |
| Total Mbases | 479.6 (54.5 – 1400) | - |
| Total Mbases (post QC) | 391.4 (59.4 – 600^) | 273 (110-404) |
| Total Reads (post QC) | 82,432 (14,790 – 167,621) | 1,215,360 (555,750 – 1,606,430) |
| Read Quality (post QC) | Q9.91 (8.8 – 11.2) | Q36 (35.28 – 36.09) |
| Read length (post QC) | 4810 bp (2691 – 10511) | 236 bp (215 – 247) |
| N50 (post QC) | 6819 bp (4031 – 11748) | - |
| **Contamination\*** |  |  |
| Per run (%) | Classified reads: (0.97-3.55)<br>Total reads: (1.10-3.75) | - |
| Per isolate (%) | Classified reads: (0.22-5.79)<br>Total reads: (0.25-6.25) | Classified reads: (0.36-4.87)<sup>^</sup><br>Total reads: (0.58-5.84) |
| **Assembly\*\*** |  |  |
| Estimated Coverage; mean (median) | 96x (84x) | 109x (113x) |
| Total Chromosomal Contigs; mean (median) | 3.5 (1) | 54.7 (51.5) |
| Total Plasmid Contigs; mean (median) | 5.4 (4) | 39.1 (38.5) |

*\*Contamination is defined as any read not classified as Escherichia (genus level).*
*\*\*Assembly after polishing and consensus. <sup>^</sup>The QC step took the top 600Mbases for assembly if there were more total Mbases than 600.*
*<sup>^</sup>Does not include contaminated isolate that was removed from subsequent analyses.*

Final polished isolate assemblies of the long-read data from the Nanopore MinION were compared to polished isolate assemblies from short-read data of the Illumina MiSeq, hybrid assemblies, and to phenotypic properties generated by AST. The concordance between long-read assemblies and hybrid assemblies was consistently higher than the concordance between short-read assemblies and hybrid assemblies for ARG identification (98% vs 88%, respectively, n=114 ARGs), identifying the genetic component – plasmid or chromosome – hosting the ARG (99% vs 66%, respectively, n=114 ARGs), identifying virulence factors (96% vs 90%, respectively, n=209 virulence factors), identifying the genetic component hosting the virulence factor (96% vs 79%, respectively, n=209 virulence factors), and serotype (100% vs 98%, respectively, n=42 isolates). See Tables S2-S5 in the supplementary material for additional detail. However, when single nucleotide polymorphism-based (SNPs) methods were used the short-read assemblies performed better (cgMLST and Prokka annotated phylogenetic trees). Phylogenetic trees demonstrated more consistent clustering between short-read assemblies and hybrid assemblies of the same isolate as compared to long-read assemblies. Next, the Tricycle Protocol and AST ARG presence/absence results were compared to the long-read assembly results. Table S6 lists the antibiotic tested and positive identification across phenotypic (AST/Tricycle) and genotypic (Nanopore) approaches. Concordance of the of the long-read assemblies with the AST and Tricycle methods was 91% (143/158 phenotypic opportunities). The AMR classes with concordances below 90% included Phenicol (3/5, 60%), Fluoroquinolones (23/28, 82%), Aminoglycosides (11/13, 85%), and Tetracycline (20/23, 87%) while concordance for B-lactamases was 98% (50/51). Finally, the bioinformatic pipelines built on Galaxy Europe for both long-reads (Nanopore) and short-reads (Illumina) were compared to previously validated ‘non-Galaxy’ pipelines (command line tools available on Github). Outputs and results suggested both pipelines were concordant between the Galaxy and non-Galaxy implementation.

### Whole Genome Sequencing: Oxford Nanopore

The sequenced isolates were first evaluated for the presence of ESBL genes. Overall, 20 of 21 (95%) of phenotypically confirmed ESBL isolates had an ESBL gene present while 13 of 21 (62%) of the general isolates (no antibiotic selection) had an ESBL gene present. Among the phenotypically confirmed ESBL isolates, a blaCTX-M gene was present on 19 of 21 (90%) isolates with one isolate having only a blaTEM gene and 9 of 21 (43%) having both a blaCTX-M and blaTEM gene present. Of the blaCTX-M genes present in the phenotypically confirmed ESBL isolates (n=21), 14% (3 of 21) were blaCTX-M-1, 14% (3 of 21) were blaCTX-M-15, and 67% (14 of 21) were blaCTX-M-55 (one was blaCTX-M-27). Among the general isolates, blaTEM was most common (12 of 21; 57%) and blaCTX-M was only present in 4 of 21 (19%) isolates (three blaCTX-M-55 and one blaCTX-M-180). In addition, 19 of 21 (90%) of phenotypically confirmed ESBL isolates carried the ESBL gene(s) on a plasmid while 4 of 21 (19%) carried the ESBL gene(s) on the chromosome (two isolates harboured ESBL gene(s) on both). For general isolates, 12 of 21 (57%) carried the ESBL gene(s) on a plasmid while 2 of 21 (10%) carried them on the chromosome (one isolate harboured ESBL gene(s) on both). See Figure S2 in the supplementary material for additional detail.

Figure 5 depicts the distribution of ESBL genes between river samples (n=12, one upstream and one downstream at each chicken slaughterhouse) and effluent samples (n=9, pre-treatment effluent for six slaughterhouses and post-treatment of three slaughterhouses that had treatment). Among phenotypically confirmed ESBL isolates, all isolate genomes contained either blaCTX-M only or blaCTX-M and blaTEM except two isolates (one isolate carried only the blaTEM gene while the other carried no specific ARG related to the ESBL phenotype). Both river and effluent sample proportions were similar between the two groupings. For the general isolates, the majority of *E. coli* isolated from river samples had no ESBL genes, while the majority of effluent samples had the blaTEM gene. However, the majority of the blaTEM genes were blaTEM-1B/C/D with one blaTEM-128 gene variant.

**Figure 5.**
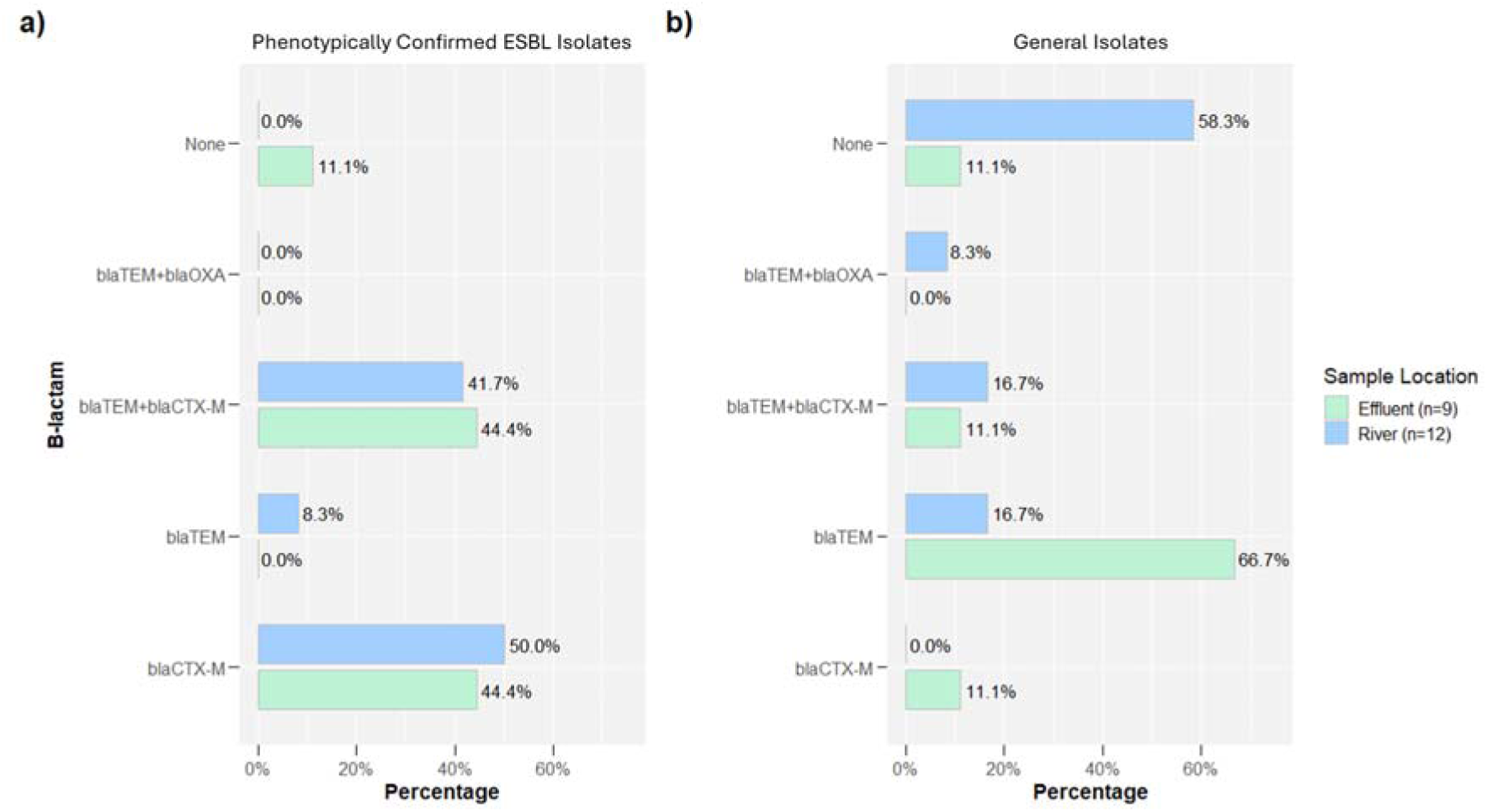
Prevalence of ESBL genes among a) phenotypically confirmed ESBL isolates (n=21) and b) general isolates (n=21). The one blaOXA gene was the blaOXA-10.

Figure 6 depicts the percentage of isolates which carried ARGs associated with unique ARG drug classes for both phenotypically confirmed ESBL isolates and general isolates. Among phenotypically confirmed ESBL isolates, only one isolate was not genotypically multidrug resistant (defined as hosting genes from two or more drug classes), while 16 of 21 (76%) isolates carried ARGs from ≥6 drug classes. Among general isolates, 8 of 21 (38%) isolates carried no ARGs, while 8 of 21 (38%) general isolates carried ARGs for ≥6 drug classes. Figure S3 disaggregates the samples by sampling location (river vs effluent). For phenotypically confirmed ESBL isolates, there was little difference between the river or effluent samples, but for the general isolates the majority of river isolates carried no ARGs while 89% of effluent samples carried ARGs for at least 4 drug classes.

**Figure 6.**
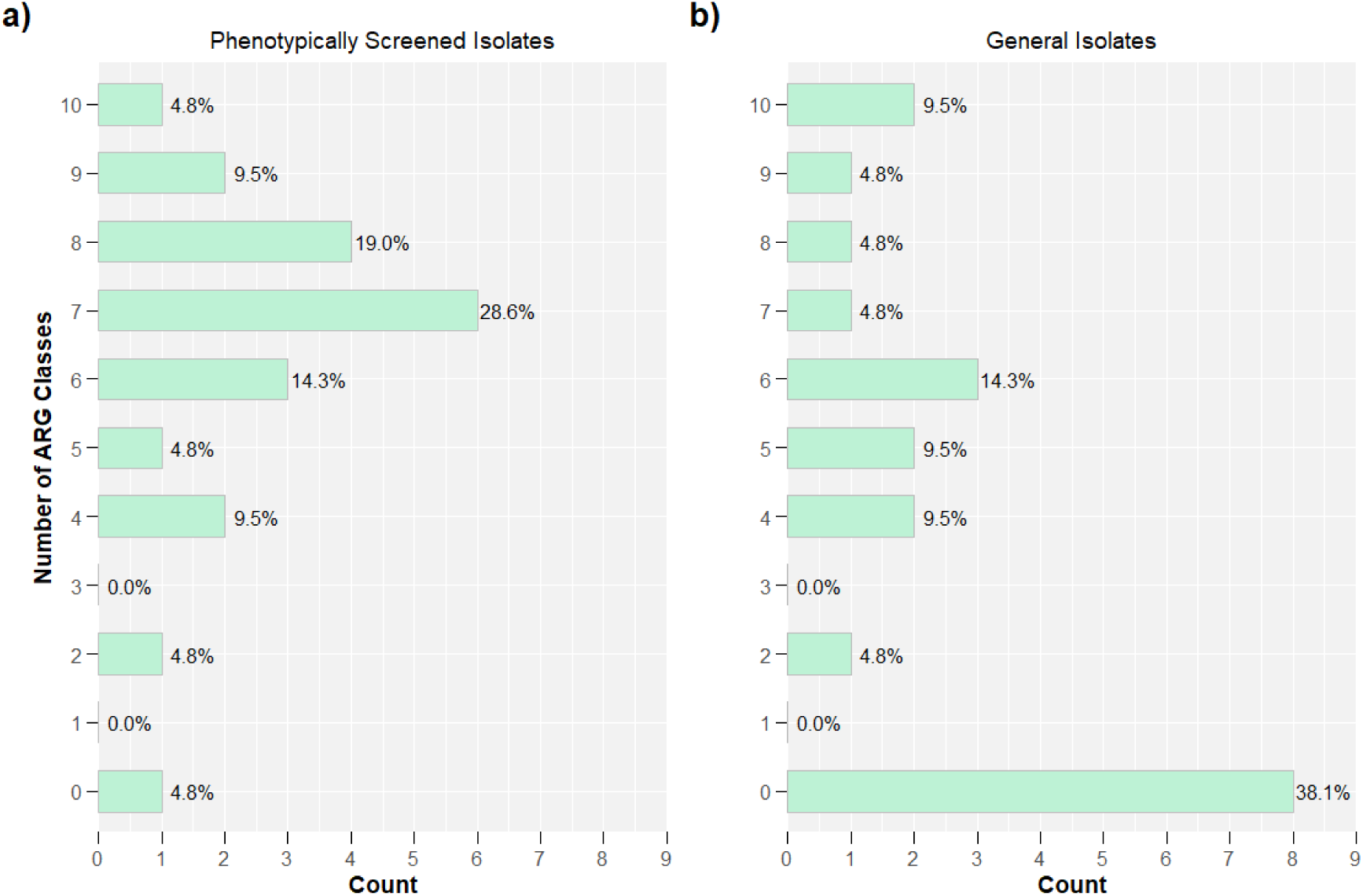
Multi-drug resistance among a) phenotypically confirmed ESBL isolates (n=21) and b) general isolates (n=21).

Among all isolates, there were 353 total ARGs, 298 (84%) carried by plasmids and 55 (16%) carried on chromosomes (see Figure S3). Of the 353 total ARGs there were 48 unique ARGs and associated variants (as denoted by ResFinder4.1). Among phenotypically confirmed ESBL isolates, there were 214 total ARGs, 171 (80%) carried by plasmids and 43 (20%) carried on chromosomes. For general isolates, there were 139 total ARGs, 127 (91%) carried by plasmids and 12 (9%) carried on chromosomes. Figure 7 and S4 present the abundance of all ARGs by phenotypically confirmed ESBL versus general isolates and by sampling location (effluent vs river). Of particular note is all Quinolone resistance genes were found on plasmids. In addition, all Phenicol resistance gene were found in the isolates not phenotypically confirmed for ESBL (i.e., general isolates). A higher proportion of effluent isolates had B-lactam resistance (ESBL), confirming Figure 5, however river samples had a higher proportion of Aminoglycoside, Phenicol, Macrolide, and Lincosamide resistance genes. In addition to B-lactam resistance genes, effluent samples also had a higher proportion of Quinolone, Sulphonamide, and Tetracycline resistance genes.

**Figure 7.**
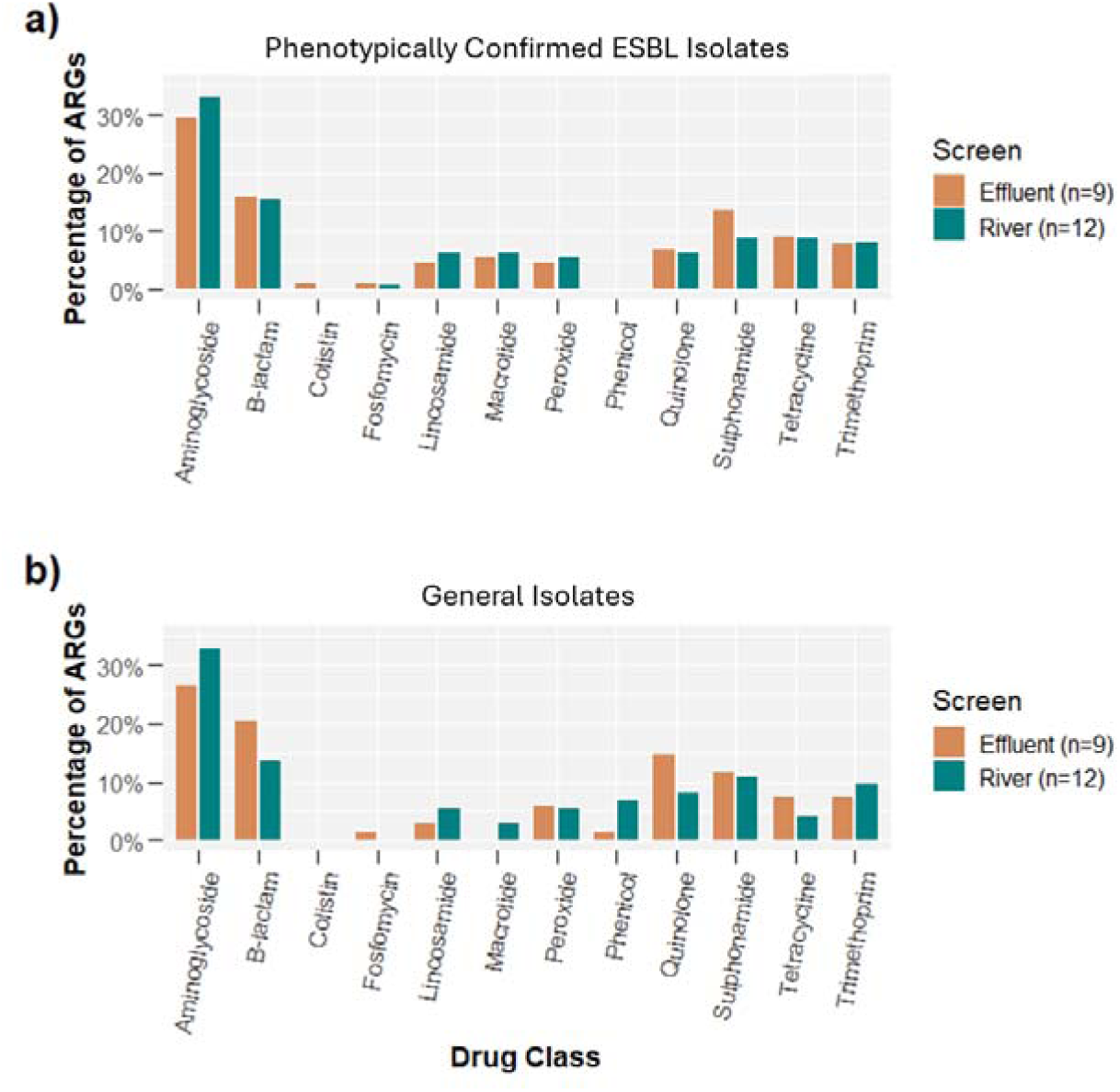
Prevalence of ARGs by drug class disaggregated by a) phenotypically confirmed ESBL isolates and b) general isolates and by sampling location.

Figure S5-S7 depict the distribution of pathotypes identified via virulence factors among both groups of isolates. For both groups, intestinal pathogenic *E. coli* (IPEC) was only identified once while extraintestinal pathogenic *E. coli* (ExPEC) had a higher prevalence. However, phenotypically confirmed ESBL isolates were classified as ExPEC more often than general isolates (62% vs 43%, respectively). Overall, of the general isolates, 11 of 21 (52%) were non-pathogenic, while among phenotypically confirmed ESBL isolates only 7 of 21 (33%) were non-pathogenic.

Finally, to evaluate relatedness between isolates, phylogenetic trees were constructed based on shared core genes (genes present in ≥95% of isolates). Figure S8 and S9 depict the phylogenetic trees for the phenotypically confirmed ESBL isolates and the general isolates, respectively. Sampling location, pathotype, and presence of ARGs and plasmids were compared to clusters within the phylogenetic tree. However, no significant trends were identified given the metadata evaluated. Further details are provided in the supplementary material.

### Cost Analysis

The cost of Nanopore sequencing will depend on the intended use within the surveillance system, the market cost of the reagents, and the ease of access to reagents. Table S7 presents the costs of reagents (as of 2019) used in this isolate-based portion of the pilot study separated into capital costs, reagent costs, and data management costs. No personnel costs are included in this initial table. Costs per sample were estimated but did not include reagents used for training sessions. Table 3 presents the total cost per sample which includes capital costs (e.g., laptop, MinIT device, hard drives) and a cost per sample excluding capital costs to provide information for extended use of this approach. Finally, we provide a cost estimate when optimal lab processing conditions are met (eight barcoded samples per run; excluding capital costs) as well as an optimal cost for isolation using the Tricycle Protocol.

**Table 3.** Costs and capabilities of approaches and technologies for isolate analyses.

|  | **Tricycle Protocol** | **MinION Sequencing** | **MiSeq Sequencing** | **Hybrid Approach** | **Sentitire AST** |
| --- | --- | --- | --- | --- | --- |
| Pilot Study Cost-per-Sample | \$10 | \$292 | \$225 | \$517 | \$20 |
| Projected Cost-per-Sample at Scale* | \$5 | \$70-100 | \$80-150 | \$150-250 | \$12 |
| ESBL E. coli Concentration | **X** |  |  |  |  |
| Total E. coli Concentration | **X** |  |  |  |  |
| ESBL to Total E. coli Ratio | **X** |  |  |  |  |
| ARG Identification |  | **X** | x | **X** | **X** |
| ARG Location |  | **X** | x | **X** |  |
| Virulence Factor Identification |  | **X** | x | **X** |  |
| Virulence Factor Location |  | **X** | x | **X** |  |
| Serotype |  | **X** | x | **X** |  |
| Phylogenetic Relatedness |  | x | **X** | **X** |  |

*Smaller 'x' denotes where a method was assessed to be less capable in our analysis.*
*\*Estimates from local suppliers in Indonesia.*

## Discussion

The first question evaluated by this pilot study was: *what types of E. coli, concentrations, and associated resistance genes are present in the effluent of chicken slaughterhouses and the receiving rivers*? First, while pathogen and ARG control is an important feature on chicken farms, the aim of the slaughterhouse is to ensure efficient processing while separating or neutralizing naturally contaminated portions of the chicken. It is this separated waste stream (i.e., the effluent) that can be monitored and potentially treated as necessary (dependent on the context). Results suggest that while both general and ESBL *E. coli* concentrations were higher in effluent, these values were within the range of values reported globally for rivers (Berendes et al., 2020; Wang et al., 2022). However, for recreational water bodies (e.g., rivers in which humans swim or play), EPA recommendations are for a geometric mean of 126 CFU/100mL for *E. coli* (US Environmental Protection Agency, 2012). The proportion of ESBL *E. coli* to general *E. coli* can be a useful indicator for ARG surveillance and data from this pilot study suggested pre-treated effluent ratios were twice as high as river ratios (median proportions: 14.2% vs 6.2%, respectively). Previous data from Wang and colleagues evaluated *E. coli* levels across types of water sources from 10 countries, with open drain and flood water having the highest levels of *E. coli* concentrations (range of means 4.38 to 7.99 log10 values) (Wang et al., 2022). However, even our limited data support the extensive body of literature on the effectiveness of treatment technologies on AMR control (Nguyen et al., 2021; Uluseker et al., 2021; Wang and Chen, 2022; Zhu et al., 2021). The two slaughterhouses which had multi-stage, fully operational effluent treatment demonstrated reductions in *E. coli* and the ratios of ESBL to general *E. coli* (general *E. coli* 6.72 log10 to 3.22 log10 and a 7.2% ratio reduction). Identifying cost-effective treatment methods adapted to local approaches, can provide critical control mechanisms for the spread of AMR.

Isolate characterization suggested that among general isolates (those with no specific AMR selection), 6 of 9 effluent isolates carried blaTEM (one had no ESBL genes), while 7 of 12 river isolates had no ESBL genes (Figure 5). Only 6 of 21 general isolates carried an ESBL gene that was not blaTEM 1B/C/D. Among phenotypically confirmed isolates, 14 of 21 blaCTX-M genes isolated were blaCTX-M-55 (similar proportions across effluent and rivers). Evaluating all drug classes among general isolates, in rivers, the most prevalent ARGs included genes conferring resistance to aminoglycosides (33%), beta-lactams (14%), and sulphonamides (11%) while in effluent genes conferring resistance to aminoglycosides (26%), beta-lactams (20%), and quinolones (14%) were most prevalent. According to FAO and DGLAHS, MoA internal survey data on antimicrobial usage in broiler poultry in 2020, enrofloxacin and amoxicillin-colistin were among the most extensively used by poultry producers in Indonesia. However, colistin use has decreased significantly since 2017. The intensive use of antibiotics in 2020 was aimed to prevent infectious diseases in broilers. This could result in *E. coli* resistance to antibiotic classes found in slaughterhouse effluents (personal communication). When isolates were phenotypically screened for ESBLs, the results of sequencing suggested the prevalence for ARGs in isolates from rivers and effluent were similar. In addition, whole genome sequencing and gene annotation demonstrated that ARGs were typically carried by plasmids. Finally, the pathotype of isolates was evaluated via the virulence factors present in the genome assemblies. Only two isolate genomes contained virulence genes associated with IPECs (both isolated from river samples). However, the majority of isolate genomes contained virulence genes associated with ExPECs (22 of 42; see Figure S4).

The second question evaluated by this pilot study was: *can the Oxford Nanopore MinION sequencing technology provide valid and cost-effective data for AMR monitoring systems currently deploying the Tricycle Protocol?* To evaluate the validity of data generated by the Nanopore MinION, outputs were compared to two ‘gold standard’ methods, namely, hybrid short-long read assemblies (following the PulseNet International protocol for Illumina MiSeq sequencing) and AST. The comparison between the long-read assemblies (i.e., MinION) and hybrid assemblies enabled evaluations of sequencing outputs and bioinformatic pipelines (using open-source, non-command line tools). Several recent studies reviewed accuracy of short-read, long-read, and hybrid assemblies identifying pro and cons to both approaches (Foster-Nyarko et al., 2022; Neal-McKinney et al., 2021; Oh et al., 2022). Recently, Foster-Nyarko and colleagues evaluated Nanopore-only performance, suggesting current methods (used in this study) are robust for key isolate characteristics, however, due to the high error rate, cluster analysis in phylogenetic trees is still limited. This was confirmed in this pilot study. Polished assemblies from the hybrid and long-read pipelines demonstrated robust concordance for ARGs (98%), virulence factors (96%), and serotypes (100%). However, the cgMLST and phylogenetic trees (using cgMLST) struggled to align. Interestingly, Foster-Nyarko and colleagues suggest a SNP limit of 50 may be necessary to rule out a long-read-only assembly from a cluster within a tree. This suggests that improved error rates or assembly correction may be needed prior to using both long-read and short-read assemblies within the same phylogenetic tree. Lastly, the ARGs identified using the long-read assemblies achieved a concordance of 91% with the phenotypic AST results. These results provide initial evidence that the Nanopore MinION can generate comparable results to gold standard genetic and phenotypic methods. Finally, the comparison between the long-read and short-read bioinformatic pipelines used here is important in two ways. First, the short-read pipeline has been available since 2013 and validated (Griswold et al., 2021), where the long-read pipeline is still being tested by various stakeholders (personal communication). Second, a key aim in this pilot study was to utilize free, open-source, and easily deployable bioinformatics tools and therefore the adaptation of both pipelines from command line-based tools to the browser-based Galaxy Europe tools and gene identification and annotation tools on the Center for Genomic Epidemiology’s website needed to be validated. Adaptation comparisons suggested differences in outputs were negligible between bioinformatic platforms used.

The Nanopore MinION can provide flexibility in national AMR surveillance systems, but also comes with specific limitations. If certain efficiencies are obtained by the laboratory, per sample costs can be around $81 or lower excluding capital costs. However, capital costs are minimal enabling more equitable access supported by the simplified library preparation (reducing lab time and need for extensive training). Comparatively, Illumina sequencing equipment is not mobile and can cost at least $100 - $225 per sample depending on operations and batching. While the MinION device is extremely portable, the reagents are less portable and often have a short shelf life. For example, both the Rapid Barcoding kit and flow cells must be stored at 2-8 °C and have a 90-day warranty (Oxford Nanopore Technologies, United Kingdom). However, experience suggests flow cells can last up to a year in proper storage. To normalize fluctuations in reagent needs in a given region, a centralized distribution channel needs to be established to help increase access and decrease the chance of expired reagents. However, supply chains and import processes can inhibit efficient and sustainable use if not addressed appropriately. Finally, the frequency of sampling monitoring points will be site specific. The World Health Organization (WHO) has recommended that for the Tricycle Method, samples from an individual site be taken monthly (World Health Organization, 2021). If monetarily feasible, all isolates should be sequenced, but an agreed upon subsample can also be used. This should be selected based on the monitoring points in a region and environmental factors, as discussed in the WHO Tricycle Method handbook. If these operational hurdles can be overcome, the additional information provided by sequencing, as demonstrated by this pilot study, can be a valuable information asset to current national AMR surveillance systems.

Limitations in this pilot study included variation in slaughterhouse characteristics, sample size and cross-sectional sampling at different time points, seasonality, and sequencing disruptions. As the study resources and timeline were limited, only six slaughterhouses were selected for evaluation via a cross-sectional study design. However, cross sectional sampling was conducted over several months (not all on one day). This can contribute to variation in results due to rain events and general seasonality, especially for the river sampling. However, the primary focus of this pilot study was on testing sampling, laboratory, and bioinformatic protocols. In addition, strict sampling criteria were used to improve the level of standardization in sampling environments (see supplementary material for details). Finally, for two isolates, Nanopore sequencing had been initiated, but was then disrupted due to power or internal processing errors. If this occurred, the system was restarted following the manufacturer’s instructions.

## Conclusion

This pilot study aimed to evaluate 1) *E. coli* from chicken slaughterhouses and 2) the feasibility of using the Oxford Nanopore MinION in national AMR surveillance systems. Results suggest that chicken slaughterhouse effluent has higher concentrations of *E. coli* and a higher abundance and diversity of ARGs as compared to receiving rivers. However, locally adapted treatment may be able to provide a critical control mechanism when appropriate. The Nanopore MinION produced similar outputs to hybrid assemblies (via Illumina MiSeq) and AST. Maintaining standardization and rigor while increasing access and options is critical for locally optimizing AMR surveillance of critical control points in one health systems.

## Acknowledgements

The authors are grateful for the hard-working field and lab teams that assisted in this study along with the staff of the Indonesian Food and Agriculture Organization and Government of Indonesia. In addition, the authors thank the United States Agency for International Development and the American people for their financial support. The funders did not have any influence on the design or interpretation of results of the study.

