## Supplemental Material for "Integrating Nanopore MinION Sequencing into National Animal Health AMR Surveillance Programs: An Indonesian Pilot Study of Chicken Slaughterhouse Effluent and Rivers"

### Sampling Protocol

- Fill out sampling control card;
- Collect 2 liters of sample from each sampling site (waste water from
- slaughterhouses and slaughter points, and river water from upstream, and downstream areas of slaughtering facility) by taking 300 mL samples 7 times with a difference of 5 minutes each time sampling. Sample should be collected from 20 – 30 cm below the water surface;
- Measure temperature, pH, and chlorin content of each sample collected (effluent, river water from upstream and downstream areas);
- Divide samples into 3 main parts: 600 mL for metagenomic sequencing with Nanopore, 300 mL for enumeration, isolation and identification of general and ESBL *E. coli* and 1000 mL as backup sample;
- Keep samples at 4 – 8 °C and submit samples to BPMSPH within maximum 5 hours after collection.

*General note:*

- For sampling at upstream area of river water, it is preferred to be done approximately 10 meters from effluent discharge point;
- For sampling at downstream area of river water, it is preferred to be done approximately 5 meters from effluent discharge point;
- Sampling time preferred is on the middle of operational period i.e., if slaughtering process is done from 8 AM to 12 AM, therefore sampling will be done at 10 AM.

### Tricycle Protocol

*Enumeration, isolation, and identification of general and ESBL E. coli from effluent and river water sample and ESBL E. coli confirmatory test:*

- Mix the sample well by swirling vigorously 25 times or by using vortex;
- Dilute samples 10-fold using sterile PBS (at 1:9 ratio) and mix the sample well;
- Discard the pipet used to make this dilution and use the new sterile pipet to remove 1 mL from this dilution to make the next dilution using the same method;
- Repeat these steps until all serial dilutions are made;
- Filter 1 and 3 mL of each diluted sample using 45 µm sterile membrane filter;
- Remove and place membrane filter onto plate containing TBX medium for enumeration of general *E. coli* and TBX-cefotaxime for ESBL, then incubate the plates at 35 to 37 °C for 18 – 24 hours;
- After the incubation, observe and enumerate the number of general and presumptive ESBL *E. coli* colony. Choose the plate that has ≤ 100 CFU for enumeration.
- Take 10 general *E. coli* colonies from one of the plates of each sample that has ≤100 CFU and streak onto MacConkey agar and 10 presumptive ESBL *E. coli* colonies

onto MacConkey-cefotaxime plate, then incubate the plates at 35 to 37 °C for 18 – 24 hours

- Transfer general *E. coli* and presumptive ESBL *E. coli* onto Nutrient Agar (NA) plate and grow overnight at 35 to 37 °C.

*Sulfide, Indole, Motility test (SIM)*

- Take single colony of organism from pure culture, make one stab down the center of the SIM medium using inoculating wire to within 2 millimeters of the bottom
- Incubate inoculated tube within 18 – 24 hrs at 35 to 37 °C.
- After incubation, examine the tube for motility which is evident by diffuse growth throughout the media. *E. coli* will produce indole positive motility result.
  Examine for $H_2S$ production which is shown by the presence of a black precipitate. *E. coli* does not reduce sulfate, therefore no black color along stab line will be found.
  Add 2 drops of Kovac reagent to the tube. A red or fuchia ring at the surface of the media indicates a positive indole reaction. *E. coli* is indole positive.

*Methyl red test, Voges-Proskauer test (MRVP)*

- Transfer bacterial culture into the two test tubes. Fill approximately 1 cm from the bottom of the tubes with bacterial culture
  For Methyl Red test, add three drops of methyl red into the first tube labelled as MR tube. Red color (+) indicates pH < 4.4, and yellow color (-) indicates pH > 6.2. *E. coli* will yield positive MR test.
- For Voges-Proskauer test, add 15 drops of ∝- naphthol and flick the tube. Add 5 drops potassium hydroxide into the second tube labelled as VP tube, and do not flick the tube. Take the transfer tube, take air, and blow it into the culture. Do this for every 5 minutes until the color change occurs (red color indicates a positive (+) result and no color change or brown or copper color indicates negative (-) result. *E. coli* will produce negative VP test.

*Citrate test*

- Inoculate Simmons Citrate Agar with overnight bacterial colony that is 18 -24 hrs old on the slant by touching the tip of inoculating wire
  Incubate inoculated tube within 18 – 24 hrs at 35 to 37 °C.
  Observe the growth of bacterial colony and the color change of the media. Visible growth on slant surface and the change of media from green to blue indicate positive (+) result r green-blue whilst visible growth and no color change indicate negative (-) result. *E. coli* is negative citrate.
- Store positive *E. coli* isolates in 10% glycerol media.

*ESBL Confirmatory Test*

- Grow maximum five presumptive ESBL *E. coli* on NA plates and incubate at 35 to 37 °C for 18 – 24 hours.
- Take 1 colony from each NA plate, streak each colony evenly onto separate Mueller Hinton agar plate. Conduct ESBL confirmatory test with double disc method (DDT).
- Screen each presumptive ESBL *E. coli* with cefotaxime (30 µg), ceftazidime (30 µg), combination disk of cefotaxime (30 µg) and clavulanic acid (10 µg), and combination

disk of ceftazidime (30 µg) and clavulanic acid (10 µg) and incubate the plates at 35 to 37 °C for 18 – 24 hours. *E. coli* isolate is confirmed to be ESBL positive when there is > 5 mm increase in inhibition zone generated when the isolates is screened with cefotaxime + clavulanic acid and with ceftazidime + clavulanic acid (double disc diffusion method (DTT) compared to inhibition zone generated with cefotaxime and ceftazidime alone.

- Store positive ESBL *E. coli* isolates in 10% glycerol media.

**AST Protocol**

- Inoculate ESBL *E. coli*, general *E. coli* and quality control strains (*Staphylococcus aureus* ATCC 29213, *Enterococcus faecalis* ATCC 29212, *E. coli* ATCC 25922, *Pseudomonas aeruginosa* ATCC 27853) on nutrient agar plates. Incubate the plates at 37° C for 18 to 24 hours.
- Take one single bacterial colony streak it on MacConkey agar plate. Incubate the plate at 37°C overnight.
- Take single colony and streak it on Mueller Hinton agar plate. Incubate the plate at 37° C for 18 to 24 hours.
- Prepare 0.5 McFarland suspension by mixing 3 to 5 pure colonies of bacterial isolate and measure the McFarland suspension with Nephelometer.
- Transfer 10 µL bacteria from McFarland suspension to 11 mL cation adjusted Mueller Hinton broth (CAMHB) medium.
- Inoculate Sensititre plate (EU Surveillance Salmonella/E.coli EUVSEC Plate) with 50 µL of bacterial suspension using Sensititre AIM Automated Inoculation System or Multi-channel pipette.
- Take one loop of bacterial suspension from positive control well and spread it on blood agar plate for direct colony count.
- Seal the Sensititre plate and incubate both the Sensititre and blood agar plates at 37° C for 18 to 24 hours.
- Read minimum inhibitory concentration (MIC) of each antibiotic using Sensititre Vizion or Manual View-Box and record the results.
- Conduct colony count on bacteria grow on blood agar plate.

**List of Antibiotics in Sensititre EU Surveillance Salmonella/E. coli EUVSEC Plate**

1. Sulfamethoxazole
2. Trimethoprim
3. Ciprofloxacin
4. Tetracycline
5. Meropenem
6. Azithromycin
7. Nalidixic Acid
8. Cefotaxime

9. Chloramphenicol
10. Tigecycline
11. Ceftazidime
12. Colistin
13. Ampicillin
14. Gentamicin

**DNA Extraction Protocol**

- Inoculate target bacteria in culture medium e.g. Nutrient Agar (NA) for cultivation of bacteria and incubate at 35 to 37 °C for 18 – 24 hours
- Take as many as possible colony of general and ESBL *E. coli* from each NA plate and place into a 500 µL sterile phosphate buffered saline (PBS) in a microcentrifuge tube and centrifuge at 13,000 x *g* for 5 min.
- Discard the supernatant using a pipette, taking care to not disrupt the pellet.
- Wash the pellet with 500 µL sterile PBS, and redo this step four times.
- Add 400 µl Fast Lysis Buffer to the bacterial pellet, tightly cap the tube, and resuspend the pellet by brief, vigorous vortexing.
- Transfer the entire mixture to a Pathogen Lysis Tube. Tightly cap the tube, secure it vertically or horizontally to a vortex adapter, and vortex at maximum speed for 10 min.
- Centrifuge the tube at 13,000 x *g* for 5 min.
- Transfer 100 µl of the supernatant to a fresh 1.5 ml microcentrifuge tube.

*DNA Quality Check (10 minutes)*

- If Nanodrop is available, test DNA concentration and purity. DNA concentration values obtained from Nanodrop should be adjusted downwards as over estimation occurs.

*General note:*

- For DNA counting with nanodrop, optimum value for 260/280 ratio is 1.8 +/-0.2 and for 260/230 ratio is 2.0-2.2 +/-0.2
- For 260/280 ratio, a ratio of ~1.8 is generally accepted as pure for DNA; a ratio of ~2.0 is generally accepted as “pure” for RNA. If the ratio is lower than 1.8 +/-0.2 it may indicate the presence of contaminant i.e., protein, phenol or other contaminants that absorb strongly at or near 280 nm
- For 260/230 ration a ratio of 2.0-2.2 is expected. If the ratio is lower than expected, it may indicate the presence of contaminants.

**Bioinformatics Workflows**

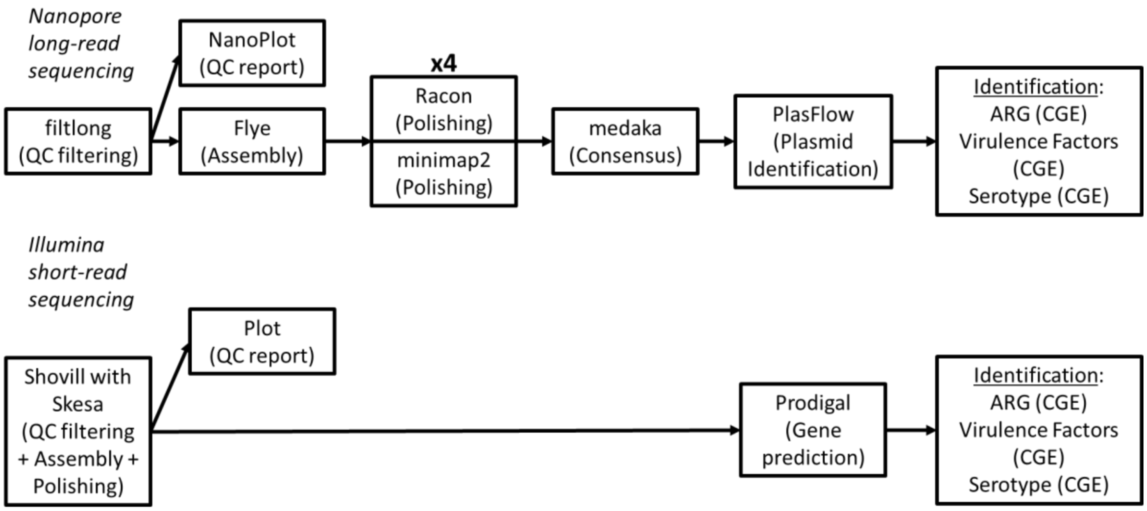

**Figure S1.** Depiction of bioinformatic pipelines

*Author's depiction. Shovill with Skesa combines several steps in the Illumina pipeline which are done individually in the Nanopore pipeline.*

### Results

#### *Nanopore Sequencing Stats*

**Table S1.** Results on QC, contamination, and assembly.

| Results per run | **Overall Mean (range)** | **Effluent (n=18); mean (range)** | **Rivers (n=24); mean (range)** |
|---|---|---|---|
| **QC** | | | |
| Total Mbases | 479.6 (54.5 – 1400) | 499.6 (74.9 – 1400) | 464.7 (54.5 – 1300) |
| Total Mbases (post QC) | 391.4 (59.4 – 600^) | 382.3 (88.7 – 600^) | 398.1 (59.4 – 600^) |
| Total Reads (post QC) | 82,432 (14,790 – 167,621) | 80,896 (23,001 – 157,777) | 83,584 (14,790 – 167,621) |
| Read Quality (post QC) | 9.91 (8.8 – 11.2) | 9.84 (8.8 – 11.1) | 9.96 (8.8 – 11.2) |
| Read length (post QC)<br>N50 (post QC) | 4810 (2691 – 10511)<br>6819 (4031 – 11748) | 4805 (2691 – 8704)<br>6724 (4031 – 11508) | 4814 (2796 – 10511)<br>6891 (4150 – 11748) |
| **Contamination*** | | | |
| Per run | Classified reads: (0.97-3.55) Total reads: (1.10-3.75) | Classified reads: (0.97-2.29) Total reads: (1.10-2.50) | Classified reads: (0.97-3.55) Total reads: (1.10-3.75) |
| Per isolate | Classified reads: (0.22-5.79) | Classified reads: (1.57-5.79) | Classified reads: (0.22-5.14) |

| | Total reads: (0.25-6.25) | Total reads: (1.85-6.25) | Total reads: (0.25-5.61) |
|---|---|---|---|
| **Assembly**** | | | |
| Estimated Coverage; mean (median) | 96 (84) | 94 (94.5) | 99 (94.5) |
| Total Chromosomal Contigs; mean (median)<br>Total Plasmid Contigs; mean (median) | 3.5 (1)<br>5.4 (4) | 3.9 (1.5)<br>6.3 (4.5) | 3.3 (1)<br>4.7 (4) |
| *Contamination is defined as any read not classified as *Escherichia* (genus level).<br>**Assembly after polishing and consensus.<br>^The QC step took the top 600 Mbases for assembly if there were more total Mbases than 600. | | | |

***Nanopore Sequencing vs Illumina Sequencing vs Hybrid Approach***

**Nanopore vs Illumina – Concordance Details**

**Table S2.** Number of ARGs identified by Nanopore, Illumina, and Hybrid Assemblies and Concordances

| No | ID isolate | Hybrid ARGs ID | Illumina ARGs ID | Illumina ARGs Located Correctly | Nanopore ARGs ID | Nanopore ARGs Located Correctly |
|---|---|---|---|---|---|---|
| 1 | AMR1_S1 | 10 | 10 | 7 | 10 | 10 |
| 2 | AMR2_S2 | 13 | 13 | 8 | 13 | 13 |
| 3 | AMR3_S3 | 0 | 0 | 0 | 0 | 0 |
| 5 | AMR5_S5 | 0 | 0 | 0 | 0 | 0 |
| 6 | AMR6_S6 | 0 | 0 | 0 | 0 | 0 |
| 7 | AMR7_S7 | 0 | 0 | 0 | 0 | 0 |
| 9 | AMR9_S9 | 8 | 7 | 7 | 8 | 8 |
| 11 | AMR11_S11 | 0 | 0 | 0 | 0 | 0 |
| 12 | AMR12_S12 | 8 | 6 | 5 | 8 | 8 |
| 13 | AMR13_S13 | 8 | 7 | 5 | 7 | 8 |
| 14 | AMR14_S14 | 17 | 12 | 11 | 17 | 17 |
| 15 | AMR15_S15 | 9 | 9 | 0 | 9 | 9 |
| 16 | AMR16_S16 | 8 | 8 | 7 | 8 | 8 |
| 17 | AMR17_S17 | 8 | 6 | 5 | 7 | 7 |
| 18 | AMR18_S18 | 0 | 0 | 0 | 0 | 0 |
| 19 | AMR19_S19 | 18 | 16 | 13 | 18 | 18 |
| 20 | AMR20_S20 | 5 | 5 | 5 | 5 | 5 |

| | | | | | | |
|---|---|---|---|---|---|---|
| 21 | AMR21_S21 | 2 | 2 | 2 | 2 | 2 |
| | Total | 114 | 101 | 75 | 112 | 113 |

**Table S3.** VFs identified by Nanopore, Illumina, and Hybrid Assemblies and Concordances

| No | ID isolate sequenced by Wates | Hybrid VF Genes ID | Illumina VF Genes ID | Illumina VF Genes Located Correctly | Nanopore VF Genes ID | Nanopore VF Genes Located Correctly |
|---|---|---|---|---|---|---|
| 1 | AMR1_S1 | 14 | 10 | 8 | 11 | 11 |
| 2 | AMR2_S2 | 11 | 9 | 8 | 10 | 10 |
| 3 | AMR3_S3 | 15 | 15 | 15 | 15 | 15 |
| 5 | AMR5_S5 | 15 | 13 | 13 | 15 | 15 |
| 6 | AMR6_S6 | 17 | 15 | 14 | 17 | 17 |
| 7 | AMR7_S7 | 6 | 6 | 6 | 6 | 6 |
| 9 | AMR9_S9 | 18 | 16 | 14 | 18 | 18 |
| 11 | AMR11_S11 | 8 | 8 | 8 | 7 | 7 |
| 12 | AMR12_S12 | 8 | 6 | 5 | 7 | 7 |
| 13 | AMR13_S13 | 8 | 7 | 5 | 6 | 6 |
| 14 | AMR14_S14 | 5 | 5 | 5 | 5 | 5 |
| 15 | AMR15_S15 | 23 | 23 | 12 | 22 | 22 |
| 16 | AMR16_S16 | 5 | 5 | 5 | 5 | 5 |
| 17 | AMR17_S17 | 6 | 4 | 4 | 6 | 6 |
| 18 | AMR18_S18 | 5 | 5 | 5 | 5 | 5 |
| 19 | AMR19_S19 | 22 | 21 | 20 | 22 | 22 |
| 20 | AMR20_S20 | 18 | 18 | 17 | 18 | 18 |
| 21 | AMR21_S21 | 5 | 3 | 2 | 5 | 5 |
| | | 209 | 189 | 166 | 200 | 200 |

**Table S4.** Comparison of Data Sets Generated by Nanopore, Illumina, and Hybrid Assemblies with SeroType

| **No** | **ID isolate sequenced by Wates** | **Hybrid O** | **Hybrid H** | **Illumina O** | **Illumina H** | **Nanopore O** | **Nanopore H** | **Illumina Match O** | **Illumina Match H** | **Nanopore Match O** | **Nanopore Match H** |
|---|---|---|---|---|---|---|---|---|---|---|---|
| 1 | AMR1_S1 | O86 | H32 | O86 | H32 | O86 | H32 | 1 | 1 | 1 | 1 |
| 2 | AMR2_S2 | O75 | H7 | O75 | H7 | O75 | H7 | 1 | 1 | 1 | 1 |
| 3 | AMR3_S3 | O83 | H1 | O83 | H1 | O83 | H1 | 1 | 1 | 1 | 1 |
| 5 | AMR5_S5 | O83 | H1 | O83 | H1 | O83 | H1 | 1 | 1 | 1 | 1 |
| 6 | AMR6_S6 | O118/O151 | H5 | O118/O151 | H5 | O118/O151 | H5 | 1 | 1 | 1 | 1 |
| 7 | AMR7_S7 | n/a | H14 | n/a | H14 | n/a | H14 | 1 | 1 | 1 | 1 |

| | | | | | | | | | | | |
|---|---|---|---|---|---|---|---|---|---|---|---|
| 9 | AMR9_S9 | O8 | H30 | O8 | H30 | O8 | H30 | 1 | 1 | 1 | 1 |
| 11 | AMR11_S11 | O178 | H20 | O178 | H20 | O178 | H20 | 1 | 1 | 1 | 1 |
| 12 | AMR12_S12 | O13/O129 | H48 | O13/O129 | H48 | O13/O129 | H48 | 1 | 1 | 1 | 1 |
| 13 | AMR13_S13 | O13/O129 | H48 | O13/O129 | H48 | O13/O129 | H48 | 1 | 1 | 1 | 1 |
| 14 | AMR14_S14 | O175 | H21 | O175 | H21 | O175 | H21 | 1 | 1 | 1 | 1 |
| 15 | AMR15_S15 | n/a | H30 | n/a | H30 | n/a | H30 | 1 | 1 | 1 | 1 |
| 16 | AMR16_S16 | O49 | H9 | O49 | H9 | O49 | H9 | 1 | 1 | 1 | 1 |
| 17 | AMR17_S17 | O13/O129 | H48 | O13/O129 | H48 | O13/O129 | H48 | 1 | 1 | 1 | 1 |
| 18 | AMR18_S18 | O154 | H4 | n/a | H4 | O154 | H4 | 0 | 1 | 1 | 1 |
| 19 | AMR19_S19 | O78 | H4 | O78 | H4 | O78 | H4 | 1 | 1 | 1 | 1 |
| 20 | AMR20_S20 | O18ac | H7 | O18ac | H7 | O18ac | H7 | 1 | 1 | 1 | 1 |
| 21 | AMR21_S21 | O82 | H45 | O82 | H45 | O82 | H45 | 1 | 1 | 1 | 1 |
| | | | | | | | | 94.4% | 100% | 100% | 100% |

**Table S5.** Comparison of Data Sets Generated by Nanopore, Illumina, and Hybrid Assemblies with cgMLST

| No | ID isolate sequenced | Hybrid cgMLST | Illumina cgMLST ID | Nano cgST | Illumina Match | Nano Match |
|---|---|---|---|---|---|---|
| 1 | AMR1_S1 | 55617 | 55617 | 1197 | 1 | 0 |
| 2 | AMR2_S2 | 31933 | 31933 | 31933 | 1 | 1 |
| 3 | AMR3_S3 | 59050 | 135035 | 1668 | 0 | 0 |
| 5 | AMR5_S5 | 59050 | 135035 | 1668 | 0 | 0 |
| 6 | AMR6_S6 | 114684 | 114684 | 114684 | 1 | 1 |
| 7 | AMR7_S7 | 2197 | 2197 | 2197 | 1 | 1 |
| 9 | AMR9_S9 | 59941 | 126068 | 59941 | 1 | 1 |
| 11 | AMR11_S11 | 37668 | 37668 | 37668 | 1 | 1 |
| 12 | AMR12_S12 | 143742 | 143742 | 124705 | 1 | 0 |
| 13 | AMR13_S13 | 143742 | 143742 | 4413 | 1 | 0 |
| 14 | AMR14_S14 | 70459 | 70459 | 70459 | 1 | 1 |
| 16 | AMR16_S16 | 27652 | 27652 | 96300 | 1 | 0 |
| 17 | AMR17_S17 | 143742 | 143742 | 15007 | 1 | 0 |
| 18 | AMR18_S18 | 137808 | 137808 | 137808 | 1 | 1 |
| 19 | AMR19_S19 | 114611 | 114611 | 114137 | 1 | 0 |
| 20 | AMR20_S20 | 142253 | 142253 | 23945 | 1 | 0 |
| 21 | AMR21_S21 | 63108 | 63108 | 63108 | 1 | 1 |
| | | | | | 83.3% | 47.1% |

### *Nanopore Sequencing vs AST*

**Table S6.** Results of the WGS/Nanopore-AST Comparison

| Antimicrobial Class | Antimicrobial | AMR Gene | Isolates with phenotypic resistance (n=38) | Isolates with phenotypic resistance and AMR gene | % of isolates w phenotypic resistance and ≥1 conferring AMR gene |
|---|---|---|---|---|---|
| Aminoglycosides | Gentamicin | aac(3)-IId | 13 | 10 | 85% (11/13) |
| | | aac(3)-IVa | | 1 | |
| B-lactamases | Ceftrazidime and Cefotaxime (also Ampicilin) | blaCTX-M-1 | 21 (+1 ceftazidime only) | 3 | 95% (21/22) |
| | | blaCTX-M-15 | | 3 | |
| | | blaCTX-M-180 | | 1 | |
| | | blaCTX-M-27 | | 1 | |
| | | blaCTX-M-55 | | 12 | |
| | | blaTEM-106 | | 1 | |
| | Ampicillin | blaOXA-10 | 29 | 1 (shared w TEM-1B) | 100% (29/29) |
| | | blaTEM-128 | | 1 | |
| | | blaTEM-1B | | 16 (2 shared w TEM-1C) | |
| | | blaTEM-1C | | 2 | |
| | | blaTEM-1D | | 3 | |
| Phenicol | Chloramphenicol | catA1 | 5 | 0 | 60% (3/5) |
| | | catB2 | | 1 (shared w cmlA1) | |
| | | cmlA1 | | 2 | |
| | | floR | | 1 | |
| *Polymyxins* | *Colistin** | *mcr-1.1* | *2* | *1* | *50% (1/2)* |
| *Fluoroquinolones* | *Ciprofloxacin** | *qnrS1* | *28* | *23 (1 shared w VC4)* | *82% (23/28)* |
| | | *qnrVC4* | | *1* | |
| Macrolide | Azithromycin | mph(A) | 12 | 12 | 100% (12/12) |
| Tetracycline | Tetracycline | tet(A) | 23 | 11 (2 shared w tet(B)) | 87% (20/23) |
| | | tet(B) | | 11 | |
| Sulphonamides/ trimethoprim | Sulfamoxazole; trimethoprim | dfrA1 | 19 (+2 sulf only; +3 trime only) | 4 | 96% (23/24) |
| | | dfrA12 | | 3 | |
| | | dfrA14 | | 11 | |
| | | dfrA15 | | 1 | |
| | | dfrA16 | | 1 | |
| | | dfrA17 | | 2 | |
| | | sul1 | | 2 (1 shared w sul2&3) | |
| | | sul2 | | 18 (3 shared w sul3) | |
| | | sul3 | | 5 | |
| Total | - | - | 31 | - | 91% (143/158) |

**Point mutations conferring ciprofloxacin or colistin resistance are not included in the ResFinder v2.1 database. Additional evaluation is needed for Nalidixic Acid and is excluded currently.*

### *Oxford Nanopore Sequencing*

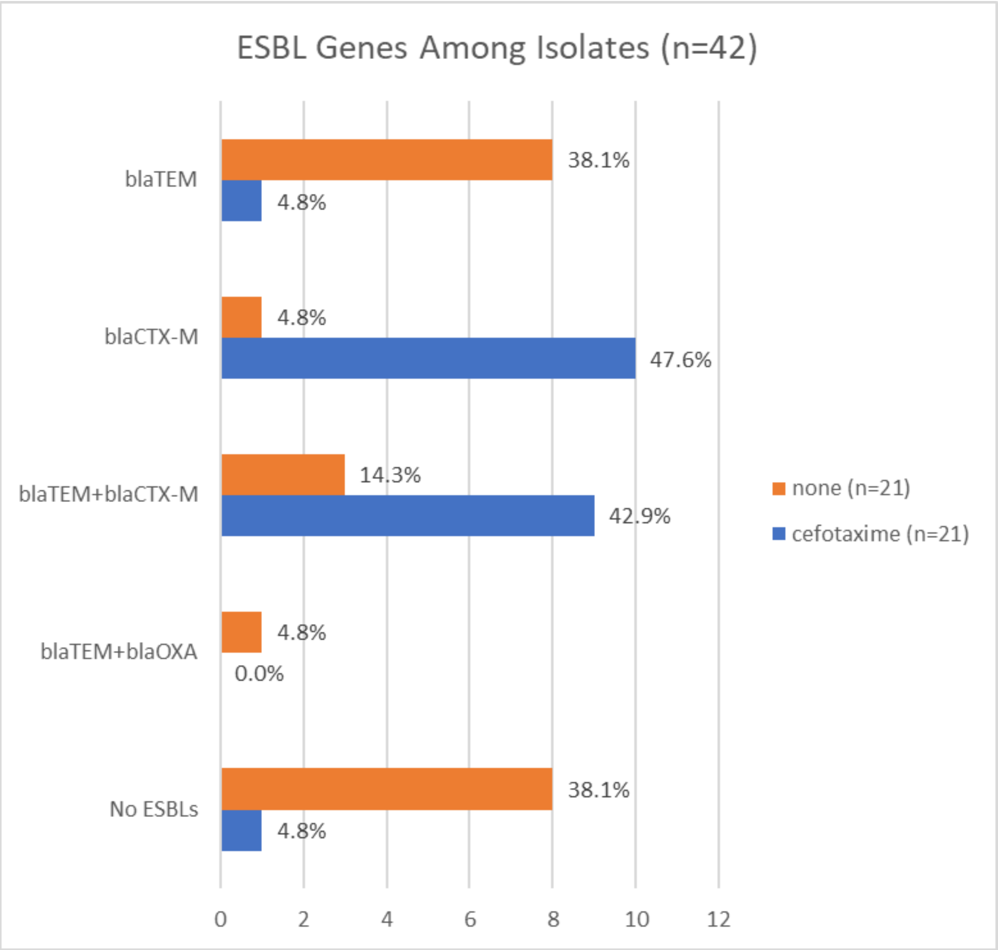

**Figure S2**. Overall distribution of ESBL genes among all samples.

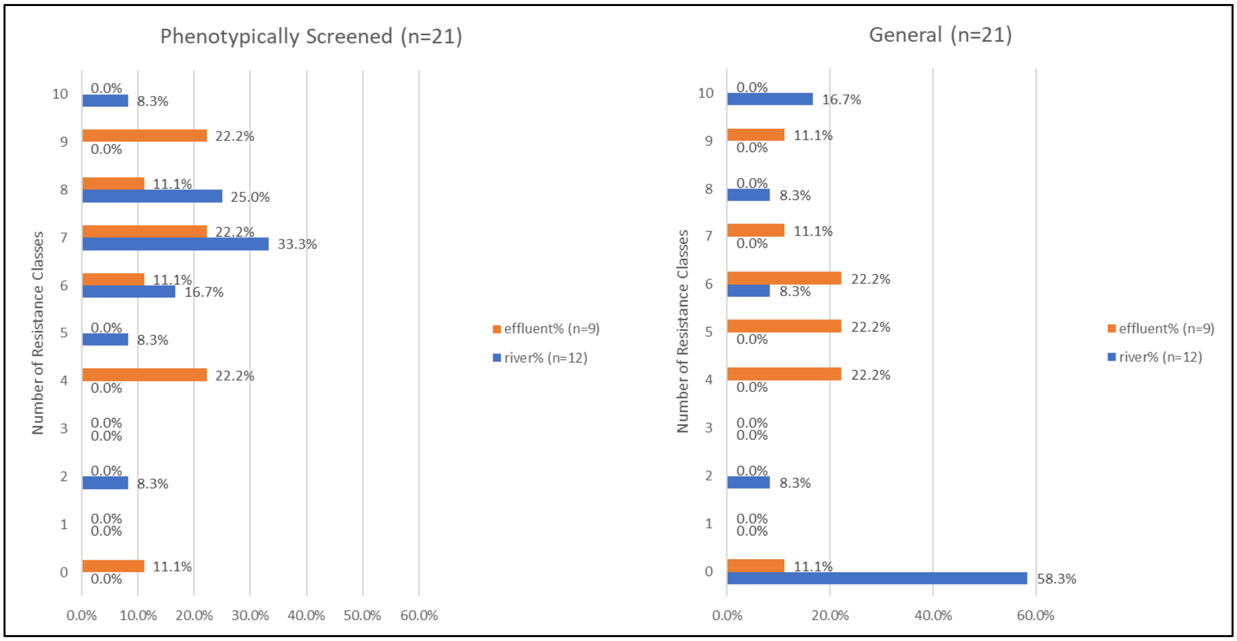

**Figure S3**. Drug resistance among isolates disaggregated by sampling location

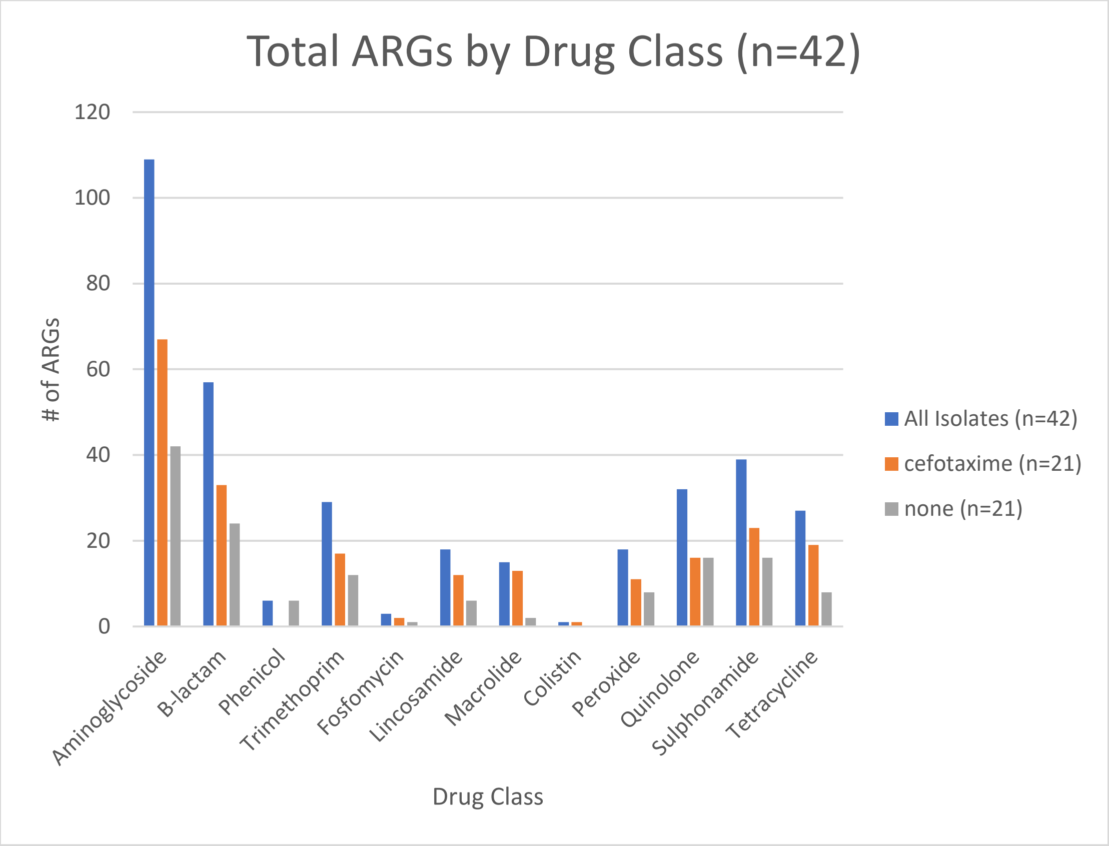

**Figure S4**. Total ARGs on all isolates

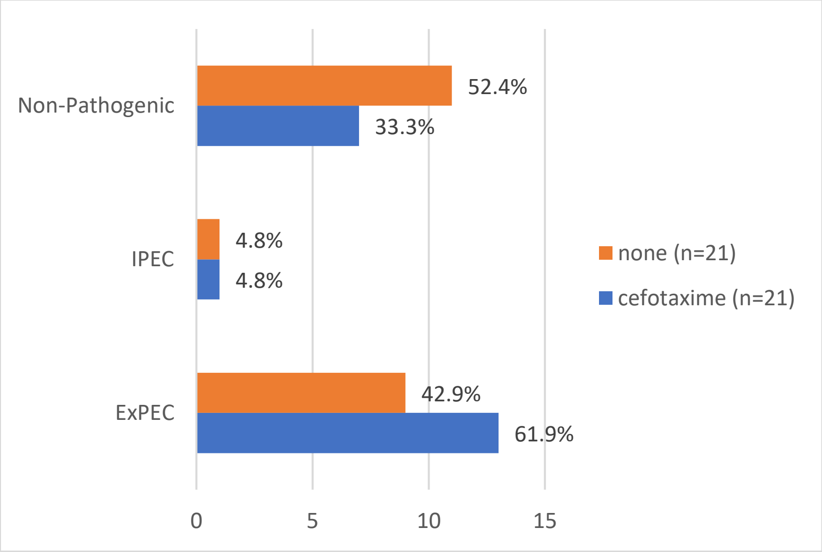

**Figure S5**. Prevalence of pathotype among all isolates

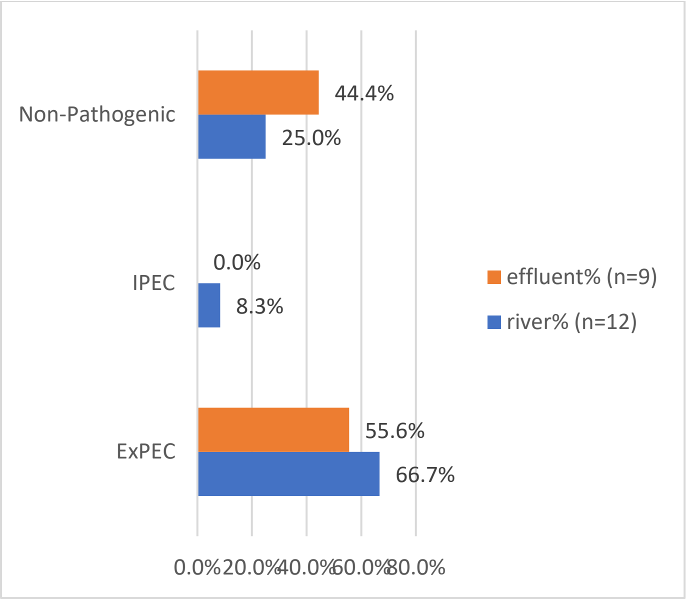

**Figure S6**. Prevalence of pathotype among phenotypically screened ESBL isolates

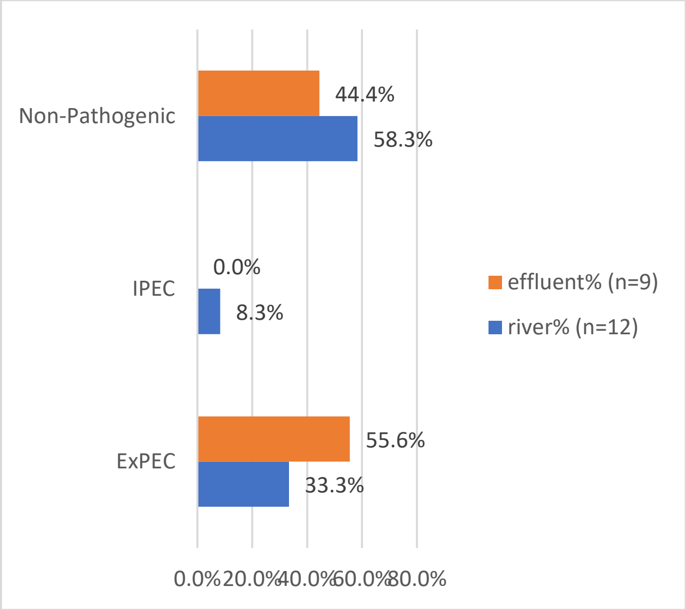

**Figure S7**. Prevalence of pathotype among general isolates.

***Phylogenetic Tree Analysis***

In Figure S8, there are three primary clusters (labelled), cluster 1 with a single isolate, cluster 2 with seven isolates, and cluster 3 with 13 isolates (which separates into two subclusters; 3a and 3b). For sampling location, the cluster 3b is primarily effluent samples while cluster 3a is primarily river samples. For pathotype, there is a visual difference between clusters 1 and 2 and cluster 3, with only 1 ExPEC present among clusters 1 and 2, while cluster 3 isolates were all ExPEC, except for 2 isolates that were not assigned a pathotype. Lastly, all but one isolate contained plasmids (as identified by PlasFlow and PlasmidFinder) and all but two isolates contained ARGs. In Figure S9, cluster 1 consisted of four isolates (the top four), cluster 2 with six isolates, and cluster 3 with 11 isolates (which separates into two subclusters; 3a and 3b).

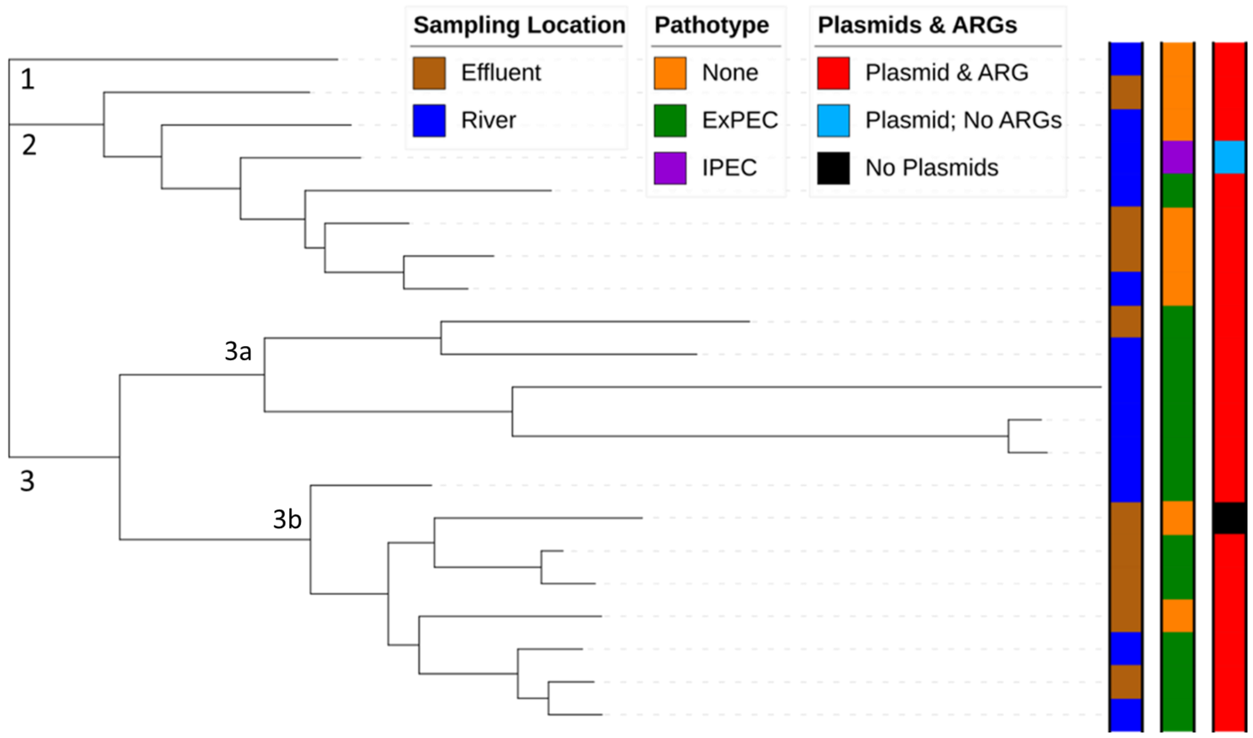

**Figure S8.** Phylogenetic tree for phenotypically screened ESBL isolates

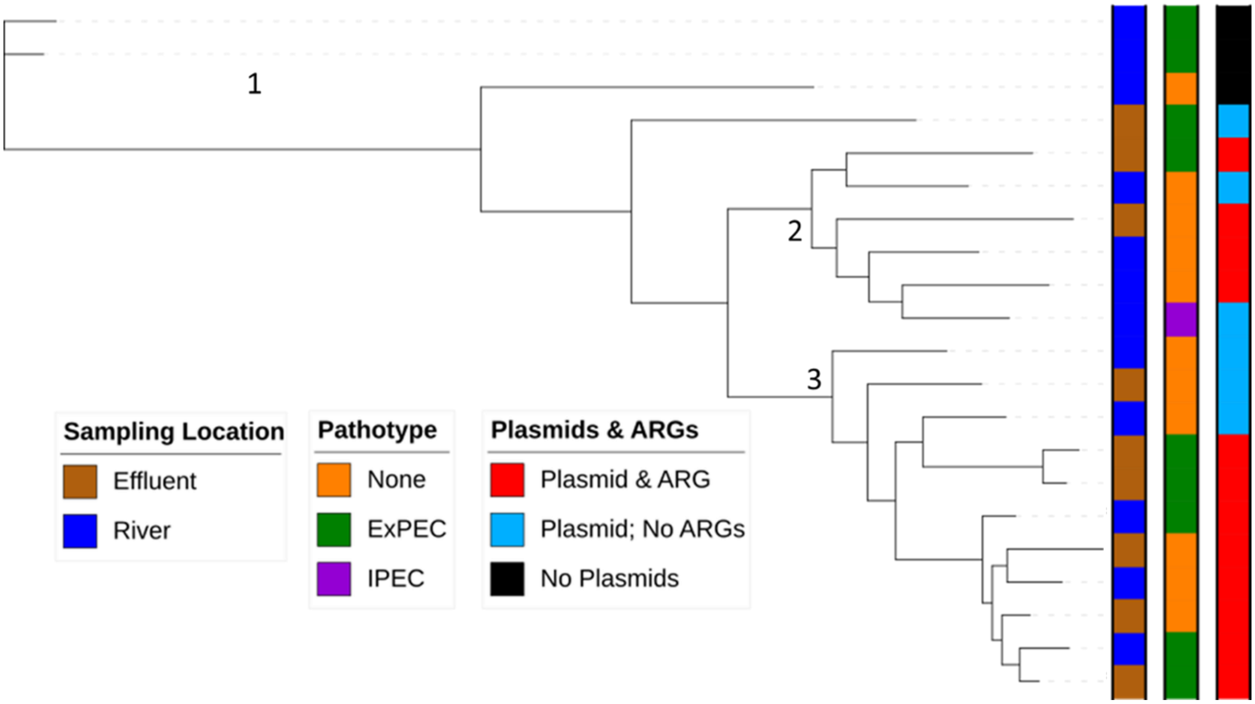

**Figure S9.** Phylogenetic tree for general isolates

***Cost Analysis***

**Table S7.** Costs of reagents used in this isolate-based portion of the pilot study.

| Item | Model # | Unit (unit price) | Total Cost |
|---|---|---|---|
| Capital Costs | | | |
| Starter Pack | | x1 | $1,000 |
| Extra MinION | | x1 | $0 |
| MinIT<br>Laptop (optional) | MNT-001 | x1<br>x1 | $2,400<br>$1,700 |
| Reagents | | | |
| 48-pack flow cells | R.9.4.1 | x1 | $24,000 |
| Rapid Barcoding Ligation Kit (6 runs/kit, 12 barcodes)<br>DNeasy PowerWater Kit (100/kit) | SQK-RPB004<br>14900-100-NF | x2 ($650)<br>x1 | $1,300<br>$974 |
| Data Management | | | |
| Data storage - Cloud (2 TB) | Google Drive | 18 months ($9.99/month) | $180 |

| | | | |
|---|---|---|---|
| Data storage – Hard drive (2TB) | Seagate portable | x1 | $70 |
| Bioinformatics tools | Galaxy Europe | - | $0 |
| Total Cost | | | $31,624 |
| **Assumes laboratory has basic supplies such as pipette tips, thermocycler (or similar), and vortex. Doesn't include sample* collection costs (drive time, collectors, etc.). | | | |
